# Hemodynamic modelling improves population receptive field estimates

**DOI:** 10.1101/2025.11.17.688931

**Authors:** D. Samuel Schwarzkopf, Ecem Altan, Steven C. Dakin, Catherine A. Morgan

## Abstract

Population receptive field (pRF) modelling is a ubiquitous tool in sensory neuroscience for estimating the functional architecture of the human brain. Most pRF models assume a canonical hemodynamic response function (HRF) to account for neurovascular effects. But how does this assumption affect results in practice? Here, we concurrently fit the HRF within the pRF model. Using simulations, we demonstrate that this algorithm accurately estimated different ground truth HRFs used for synthesizing data. Moreover, concurrent fitting improves the accuracy of pRF estimates, especially for the complex Difference-of-Gaussian model. Next, we reanalyzed empirical datasets with different stimulus paradigms and scanning parameters. Concurrent fitting substantially reduced the proportion of implausibly small pRFs. Importantly, the best-fitting HRF differed substantially from canonical HRFs or from those measured independently. HRFs also peaked earlier in higher than early visual cortex, suggesting response nonlinearities. This variability need not only be hemodynamic but could result from neural factors. Critically, all these differences could produce spurious pRF estimates and alter the interpretation of reported findings.

## INTRODUCTION

Population receptive field (pRF) modelling is a popular method for retinotopic mapping with functional magnetic resonance imaging (fMRI). Unlike the travelling wave analysis of earlier retinotopic mapping studies (Engel et al., 1997; Sereno et al., 1995; Wandell et al., 2007), pRF analysis aims to estimate the aggregated spatial response profile of neuronal populations contained in each fMRI voxel. A typical voxel in visual cortex responds only to a limited portion of visual space – analogous to the receptive field of a single neuron. Retinotopic map organization is therefore defined not only by location preferences of neuronal populations within each voxel, but also their spatial selectivity – the population receptive field size. Early visual regions typically have small pRFs while higher-level regions like the lateral occipital (LO) or middle temporal areas (MT) are much larger, sometimes covering large parts of the visual field (Amano et al., 2009; Dumoulin & Wandell, 2008). However, pRF size is not only determined by the selectivity of the underlying neuronal population, but other factors like extra-classical receptive field shapes, the distribution of neuronal receptive fields across space, and – because fMRI infers neuronal activity only indirectly by relative changes in blood oxygen concentration – it also reflects metabolic and neurovascular effects.

To estimate the pRF, most studies use the forward-modelling framework introduced almost two decades ago (Dumoulin & Wandell, 2008). This approach uses the known time series of the stimulus presented to the participant to predict the fMRI response time series in each voxel (Figure 1). Each possible pRF, which can vary in position, size, and shape, will respond differently to a given stimulus. Accurate modelling necessitates assumptions about the hemodynamic response function (HRF), which is convolved with the predicted neural response (Figure 1D-G).

**Figure 1.**
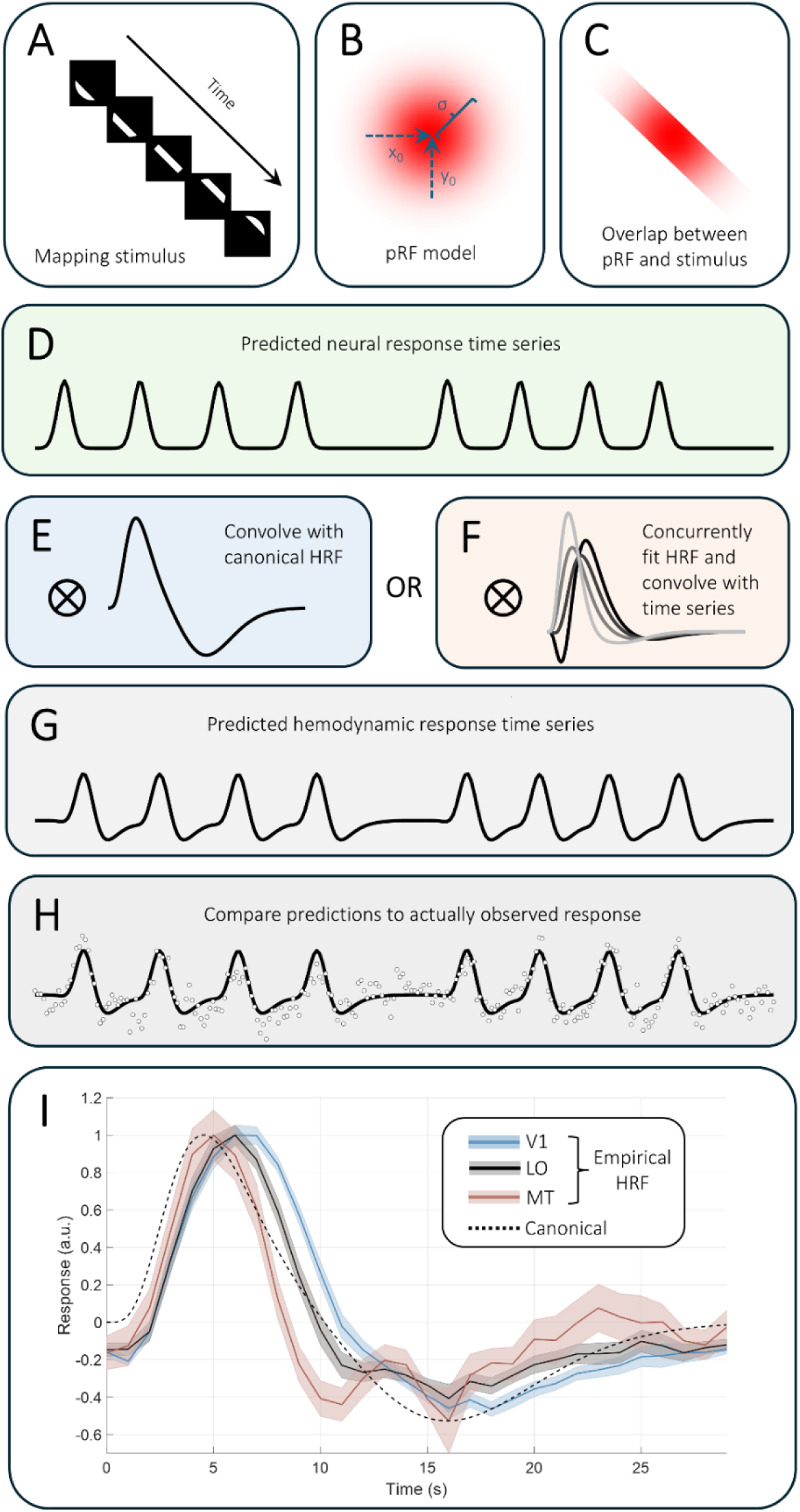
pRF modelling with fMRI. **A.** The analysis starts with knowledge of the stimulus presented to the participant during the scan. **B.** The pRF model has free parameters describing the position, shape, and response properties of the neuronal population in a voxel. The most typical pRF model is a 2D Gaussian defined by x0 and y0 position and the pRF size (σ). **C.** The amount of overlap between the pRF model and the stimulus mask predicts the neuronal response of the voxel over time (**D**). **E-G.** The predicted neural time series is convolved with an HRF. Usually, this is a canonical HRF (**E**) but in our new analysis (**F**) we fit the HRF concurrently with the pRF parameters. which then yields the predicted hemodynamic time series (**G**). **H.** The analysis optimizes the parameters of the pRF model (and, in the Concurrent fit, the HRF parameters) to achieve the closest fit between the predicted hemodynamic time series (solid black line) and the observed fMRI data (white discs). **I.** Empirically measured impulse-response functions averaged separately for areas V1, LO, and MT, and then averaged across participants (shaded regions denote ±1 standard error of the mean across participants). The dashed black line denotes the SamSrf de Haas canonical HRF for comparison. The peak latency evidently differs between these curves (V1: 6.5 s, LO: 6 s, MT: 5 s, canonical: 4.5 s).

Most pRF studies use canonical HRFs based on normative data (Boynton et al., 1996; de Haas et al., 2014; Friston et al., 1998) or individual HRFs estimated via independent, slow event-related stimulation paradigms (Dumoulin & Wandell, 2008; Schwarzkopf et al., 2014; van Dijk et al., 2016). Simulations in early studies suggested that variation in HRF shape and latency had only a negligible impact on pRF estimates (Dumoulin & Wandell, 2008). We also found that a fixed canonical HRF did not dramatically affect the reliability of pRF estimates compared to using empirical HRF fits (van Dijk et al., 2016). Since then, however, more detailed analyses showed that assuming an inappropriate HRF can strongly skew results (Lerma-Usabiaga et al., 2020), suggesting that earlier simulations failed to fully account for hemodynamic confounds. HRFs have been shown to vary substantially between participants (Aguirre et al., 1998), but also between brain regions (Handwerker et al., 2004). It also remains unclear whether HRFs estimated as impulse-response functions are appropriate for modelling the temporal dynamics of retinotopic mapping experiment, which entails stimuli moving systematically through the visual field. Because the hemodynamic response is highly nonlinear, including being dependent on stimulus duration (Lindquist et al., 2009), these impulse response functions may therefore fail to account for the actual response pattern.

Because in pRF mapping stimulus location varies with time, the HRF is effectively a spatial-temporal kernel blurring the neural response time series. It is thus inherently linked to spatial parameters like pRF size. Some researchers have used an iterative approach to fitting an HRF after first fitting a pRF model based on a canonical HRF and then refitting the model (Chang et al., 2025; de Haas et al., 2021; Harvey & Dumoulin, 2011). However, it remains unknown if this could result in the model fit becoming stuck in local minima, which could bias the estimates. More importantly, all previous studies mentioned assumed a constant HRF across the brain. This assumption could be refined in light of empirical data demonstrating that the shape of the response can vary between brain regions, in terms of its latency, duration, amplitude, and the degree to which a negative undershoot follows the initial positive response (Figure 1I). Few studies have fit the HRF separately for different brain regions (Aqil et al., 2024, 2025). Here, we therefore sought to test how fitting the HRF concurrently with the pRF model parameters impacts the results of pRF mapping studies, using both ground truth simulation and reanalysis of existing empirical datasets. To our knowledge, no study has systematically investigated how much differences in the HRF affect pRF estimates.

## MATERIALS AND METHODS

We used our custom MATLAB toolbox SamSrf (version 10) for pRF analysis (https://osf.io/2rgsm). The general pRF fitting procedures have been described previously (Altan, Boyaci, et al., 2025; Morgan & Schwarzkopf, 2019; Moutsiana et al., 2016; Schwarzkopf et al., 2014; van Dijk et al., 2016). Briefly, the model predicts the neural response of a given pRF from its overlap with the stimulus at each time point. This time series is then convolved with a given HRF. The model finds the combination of pRF parameters that maximize the fit between model prediction and observed data (Figure 1).

In most of our earlier studies, we used a canonical HRF based on an average of 28 participants individually-measured HRFs based on the impulse response to a full-screen stimulus (de Haas et al., 2014). Our novel adaptation to the pRF modelling algorithm in the current work is to estimate the HRF concurrently while estimating the pRF parameters in the fine-fitting (parameter optimization) phase. Henceforth, we refer to fitting the HRF concurrently with the pRF parameters as the *Concurrent* fit.

Specifically, we first conducted a coarse-fit (grid search) step, generating predicted time series, using a set of plausible combinations of pRF parameters (see details below). In our standard analysis, we convolved these predictions with the de Haas canonical HRF (de Haas et al., 2014). For concurrent HRF fitting, the coarse fit predictions were convolved with the classic canonical HRF used in SPM (Friston et al., 1998). We then identified the predictions in this search grid that correlated most strongly with the observed response time series for each vertex in the cortical surface data mesh. We used the parameters from this prediction to initialize parameters for fine-fitting, using the Nelder-Mead optimization algorithm (Lagarias et al., 1998; Nelder & Mead, 1965) to estimate the full model parameters.

We fit the standard two-dimensional Gaussian model, describing the pRF position (x_0_ and y_0_) and pRF size (σ) values. In our simulation analyses, we also fit an antagonistic difference-of-Gaussian (DoG) model with an excitatory center (σ_1_) and an inhibitory surround (σ_2_) size, and a parameter (δ) for the ratio of the amplitudes of the two Gaussians (Anderson et al., 2017; Harvey et al., 2013; Schwarzkopf et al., 2014; Zuiderbaan et al., 2012).

Novelly in this work, during the fine-fitting stage for the concurrent HRF fitting we also estimated five free parameters to describe the HRF. We initialized these parameters using their values from the SPM canonical HRF and constrained the range of plausible model fits: response latency = 6 s (constrained between 3-9 s), undershoot latency = 16 s (10-18 s), response dispersion = 1 s (0.5-3 s), undershoot dispersion = 1 (0.5-3 s), and response/undershoot ratio = 6 (0-10).

### Simulated data

To test the capacity of the concurrent HRF fitting algorithm, we first conducted pRF analyses using simulated data. For this, we used the standard sweeping bar stimulus from our previous work (Morgan & Schwarzkopf, 2019; Tangtartharakul et al., 2023). Specifically, we generated data for a set of pRFs with known positions and sizes, convolved these with one of three canonical HRFs (Figure 2), and then added noise from a Gaussian distribution to simulate real data. In all the analyses presented here, we used a noise with a standard deviation of 20%, which we found corresponds to a biologically plausible distribution of model fits compared to our previous empirical analyses.

**Figure 2.**
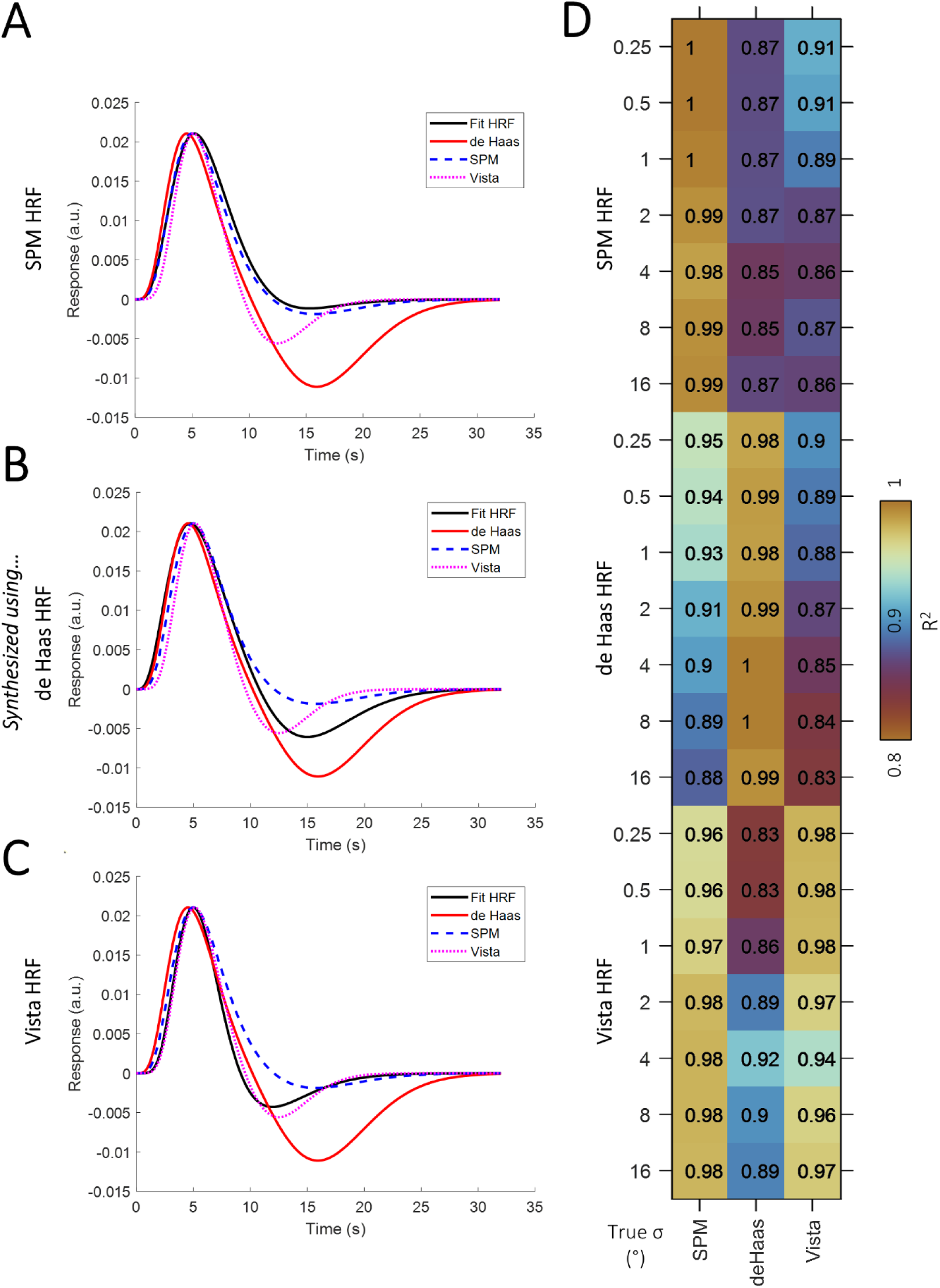
Concurrent fit closely matches the true HRF at TR=1. We synthesized data for a set of biologically plausible pRFs using either the SPM canonical HRF (**A**), our SamSrf de Haas HRF (**B**), or the Vista HRF (**C**). The black line shows the average concurrent HRF fit for each dataset, compared to the three canonical functions (see legend). **D.** Goodness-of-fit (R^2^) between the average HRF fit for each true pRF size (rows) in each of the three simulated datasets. The three columns show the R^2^ agreement between the data and fit to each of the three canonical HRFs.

For simulating two-dimensional Gaussian pRF data, we simulated ground truths at every combination of 24 polar angles (in steps of 15°), six eccentricities (0.5°, 1°, 2°, 4°, 8°, 16°), and seven pRF sizes (0.25°, 0.5°, 1°, 2°, 4°, 8°, 16°). To save computation time, for simulating the DoG pRF data, we limited the analysis to only 45° polar angle, three eccentricities (1°, 2°, 4°), four values each for σ_1_ (0.5°, 1°, 2°, 4°) and σ_2_ (1°, 2°, 4°, 8°), and three values for δ (0.125, 0.25, 0.5). We synthesized the time series for each combination of parameters twenty times and then perturbed each of them with independent Gaussian noise.

These data were then fed into our two different pRF analysis pipelines, either assuming the de Haas canonical HRF (*Canonical* fit), or with concurrent HRF fitting (*Concurrent* fit). To make the results comparable between the Gaussian and DoG models, we used identical search grids for the coarse fit of the position and the first pRF size parameter (σ or σ_1_, respectively). There were ten polar angles (in steps of 36°), nine eccentricities from 0.5° to 8° (in equal logarithmic steps), and six pRF sizes from 0.5° to 16° (in equal logarithmic steps). For the DoG model, we used the same values also for the inhibitory pRF size (σ_2_), and four values for δ (0, 0.1, 0.2, 0.3). This is different from how we have conducted the DoG analyses in previous studies (Anderson et al., 2017; Schwarzkopf et al., 2014), where we used the standard 2D Gaussian pRF model for the coarse-fit and then estimated the DoG model at the fine-fitting stage only. To note, the search grid used here was also coarser than used in previous studies and for our empirical data analysis (see below). The reason for this is that a grid search with a higher resolution of the parameter space becomes computationally impractical.

### Empirical datasets

We used several existing retinotopic mapping data from previous studies to evaluate the performance of our concurrent HRF-fitting algorithm. These analyses only used the two-dimensional Gaussian pRF model. We used a finer search grid for the coarse-fitting stage of real data than when fitting simulated data. Specifically, there were 36 polar angles, 29 eccentricities, and 34 pRF size values, with eccentricity and pRF size incrementing on a logarithmic scale.

All data were analyzed in cortical surface space. All fMRI data had been projected to the cortical surface mesh of the white-grey matter boundary generated using FreeSurfer (Dale et al., 1999; Fischl et al., 1999), using the native surface models for each participant. For the first three datasets below, visual regions of interest were determined based on a probabilistic atlas procedure (Sereno et al., 2022) and combining some of the small regions into larger clusters. Specifically, in these datasets, we analyzed data from V1, V2, V3, V4, VO1, V3A, V3B, LO, MT+, and IPS0. For the spatial frequency adaptation dataset, regions of interests (V1, V2, V3, V4, and V3A) were delineated with the automatic delineation tool of the SamSrf toolbox and then manually corrected.

#### HCP dataset

We analyzed 32 participants selected from the 7 Tesla retinotopic mapping data in the Human Connectome Project (HCP). The full dataset contains 181 participants (Benson et al., 2018); however, we selected this smaller subset, both due to computational constraints and to ensure high quality mapping data. Specifically, we limited ourselves to only those participants who showed robust and contiguous clusters of visual activation (noise ceiling > 0.3) covering occipital, ventral, parietal and prefrontal regions. The dataset contains two runs collected with an identical bar stimulus that smoothly moved through the visual field along eight directions. The run started with a 16 s blank screen epoch, another 16 s blank epoch after the fourth bar sweep, and it ended with a 20 s blank epoch. The other bar sweeps were interspersed with 4 s blank screen epochs. The carrier stimulus comprised natural images. The maximum eccentricity of the stimulus was 8°. Voxel size was 1.6 mm isotropic and the TR was 1 s, using multiband acquisition (Breuer et al., 2005). Time series for each vertex in the cortical surface mesh were detrended, z-standardized, and averaged across runs. Having an equal number of identical runs enabled us to calculate the noise ceiling from the test-retest correlations (Morgan & Schwarzkopf, 2019). We limited our data analysis to all those vertices in the cortical mesh where the noise ceiling exceeded 0.2.

#### Auckland dataset

We also reanalyzed data from 23 participants collected on a Siemens Skyra 3 Tesla scanner at the Centre for Advanced Magnetic Resonance Imaging (CAMRI) at the University of Auckland, New Zealand (Tangtartharakul et al., 2023). Voxel size was 2.3 mm isotropic and repetition time (TR) was 1 s, using multiband acquisition (Breuer et al., 2005). The study used a flickering bar stimulus containing a geometric “ripple” pattern as carrier. The bar stepped in time with each TR through the visual field along the eight directions separated by 45°. Each sweep of the bar lasted 25 s, and blank screen epochs of the same duration were interspersed after the fourth and eighth sweep of the bar. The maximum stimulus eccentricity was 9.5°. We collected six identical runs per participant. Time series for each vertex in the cortical surface mesh were detrended, z-standardized, and averaged across runs. We limited all data analysis to an occipital-temporal region of interest.

#### London dataset

The third dataset included 17 participants and was collected on a Siemens Avanto 1.5 Tesla scanner at the Birkbeck-University College London Centre for Neuroimaging (BUCNI) in London, United Kingdom (van Dijk et al., 2016). Voxel size was 2.3 mm isotropic and TR was 1 s, using multiband acquisition (Breuer et al., 2005). This study used a combined wedge and ring stimulus apertures (Alvarez et al., 2015) with natural images as carrier pattern (Moutsiana et al., 2016). Wedge rotation direction and ring contraction/expansion alternated between runs. There were three cycles of wedge rotation and five cycles of ring expansion/contraction per run. Each run ended with blank screen epochs of 45 s to estimate baseline responses. In total, there were 12 runs, split across two scanning sessions. Time series for each vertex for each stimulus direction in the cortical surface mesh were detrended, z-standardized, and averaged across runs. Then we concatenated these averaged time series from the two stimulus directions. We limited all data analysis to an occipital-temporal region of interest.

#### Spatial frequency adaptation dataset

The final dataset we analyzed was originally published as part of a study demonstrating the effect of adapting to high vs. low spatial frequency patterns on pRF size estimates in visual cortex (Altan, Morgan, et al., 2025). These data included 8 participants and were collected under identical conditions as the Auckland dataset, on the same scanner with the same stimulus sequence, with the following exceptions: before each run, participants viewed a high (3.5 cpd) or low (0.5 cpd) spatial frequency adaptation pattern (bandpass filtered white noise) for 30 s flickering at 20Hz. Subsequently, each TR contained another 500 ms presentation of the adaptation pattern, immediately followed either by a 500 ms bar stimulus or a blank interval. The bar stimulus contained a bandpass filtered white noise pattern at an intermediate (geometric mean) spatial frequency (∼1.3 cpd) between the high and low spatial frequency adapter. All runs began and ended with 25 s blank epochs during which only the fixation target was being presented.

## RESULTS

We extended our standard pRF modelling pipeline in SamSrf (https://osf.io/2rgsm) to fit the commonly used double-gamma function (Boynton et al., 1996; Friston et al., 1998) with five free parameters: the latencies and dispersions, respectively, of the initial positive response and the subsequent negative undershoot, plus a parameter denoting the ratio of amplitudes of these two components.

First, we validated this *Concurrent* fit by testing that it accurately approximates the underlying HRF used to generate synthetic data. We evaluated how this approach compares to assuming a fixed canonical HRF (henceforth referred to as *Canonical* fit) for estimating known ground truth pRF parameters. Then, we applied this algorithm to several already existing datasets collected at different magnetic field strengths (1.5, 3, and 7 Tesla) and using different pRF stimulus paradigms.

### Close match between concurrent fit and simulated HRF

A critical prerequisite for evaluating how the Concurrent fit affects pRF results is that this algorithm is in fact accurate. We first simulated time series data for a set of biologically plausible pRFs (using combinations of x_0_, y_0_, and σ parameters) and added Gaussian noise to mimic experimental data. We simulated separate datasets using three canonical HRFs commonly used in the literature (Figure 2A-C), specifically the SPM HRF (Friston et al., 1998), the de Haas HRF used until recently as default in SamSrf (de Haas et al., 2014, p. 201), and the Vista HRF (Lerma-Usabiaga et al., 2020). Each combination of ground truth pRF parameters and HRFs was synthesized 20 times to estimate the robustness of model fits in the presence of noise.

We selected all ground truth pRFs in the analysis where the goodness-of-fit of the pRF model exceeded a threshold of R^2^>0.05. This constitutes a relatively liberal threshold compared to what is used in many pRF studies, because we wanted to avoid confounding the results by only selecting good model fits. We then averaged the HRF fits for these data. We found that for stimulus designs with temporal resolution of 1 s (as determined by the repetition time, TR, of the fMRI pulse sequence), the Concurrent fit closely matched the HRF used to simulate data. Across all ground truth pRFs, the average HRF fit closely matched (all R^2^>0.94) the true HRF used to synthesize the data (Figure 2). Note that the HRF underlying the Concurrent fit is from the same family, the double gamma function, as those used the simulate data. The fit was generally best for data synthesized using the SPM HRF (Figure 2A; note that the algorithm used the parameters of the SPM HRF to initialize the parameter optimization stage). For data synthesized using the other two canonical functions, the initial response peak and shape were generally a close match, but the undershoot was somewhat misestimated; especially for the de Haas HRF the depth of undershoot was underestimated (Figure 2B). Nevertheless, the model generally performed well in estimating the simulated HRF (Figure 2D and Supplementary Figure S1). These analyses used the classic two-dimensional Gaussian pRF model. We found similar results when we instead fit Difference-of-Gaussians (DoG) center-surround models even though the true pRFs were Gaussian (Supplementary Figure S2).

The accuracy in estimating the HRF will also likely depend on the pRF parameters, especially the pRF size. We therefore also repeated this analysis with combinations of different ground truth pRF sizes. These analyses also showed close alignment between the average HRF fit and the true HRF (Supplementary Figures S1-S3). For data synthesized with the Vista HRF (Figure 2C), the fits were generally similar for the SPM and the Vista HRF; in fact, for larger pRFs the SPM HRF yielded a slightly better fit. As explained, we deliberately chose a liberal threshold for these analyses. It would be very unlikely that bad fits improve the correspondence between the estimated HRF and the ground truth. However, to check if the choice of threshold mattered, we repeated this analysis with more conservative R^2^ thresholds. If anything, this subtly improves the HRF fits to the ground truth (Supplementary Figure S4).

To note, for experimental designs with coarser temporal resolutions (TR≥2 s) the analysis yielded much worse estimates of the true HRF, especially the Vista HRF (Supplementary Figure S1). However, shorter TRs have become commonplace in pRF studies (Benson et al., 2018; Chang et al., 2025; Harvey & Dumoulin, 2011; Himmelberg et al., 2022; Morgan & Schwarzkopf, 2019; Moutsiana et al., 2016; Tangtartharakul et al., 2023; van Dijk et al., 2016). In fact, in our own group we have exclusively used a TR of 1 s in the past decade. In the following, we therefore restrict our analyses to our standard stimulus design using 1 s TR (Morgan & Schwarzkopf, 2019; Tangtartharakul et al., 2023).

### Smaller errors in pRF size estimates

We next assessed how the Concurrent fit algorithm affected the accuracy of pRF parameter estimates compared to the Canonical fit (Figure 3). In the latter, we used the de Haas HRF, because this was the canonical function used in most of our previous pRF studies using SamSrf, and its shape is distinct from the other canonical functions. However, qualitatively similar results were observed when assuming a fixed SPM HRF (data not shown).

**Figure 3.**
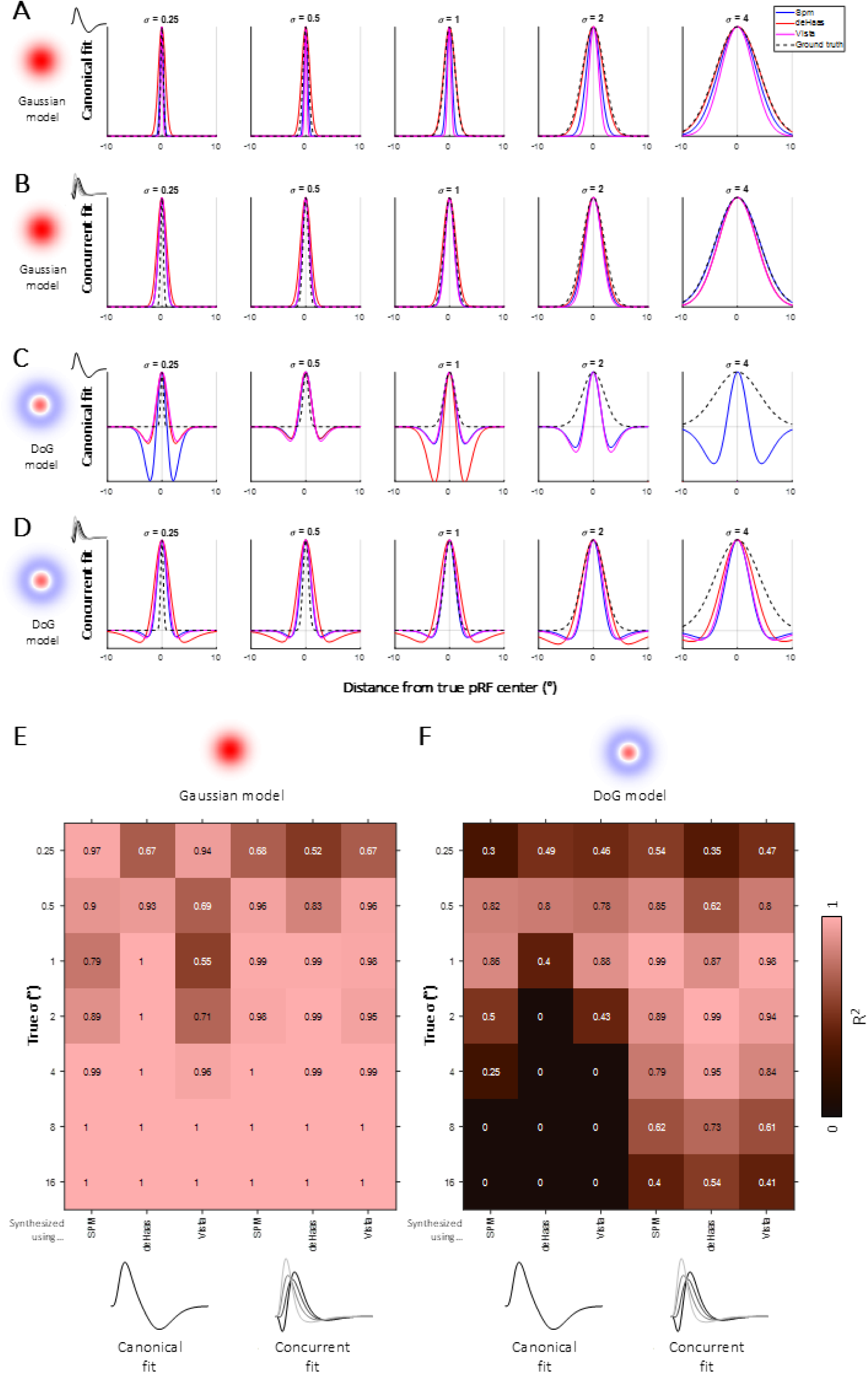
pRF estimates vs. ground truth. **A-D**. One-dimensional cross sections through the true pRF centers of simulated pRFs. The different curves depict the ground truth pRF (dashed black lines) and those estimated for data synthesized using the SPM HRF (blue), SamSrf de Haas HRF (red), or the Vista HRF (magenta). Columns show data for different ground truth pRF sizes. Results are shown for the Canonical fit (**A,C**) or the Concurrent fit (**B,D**). We fit either a 2D Gaussian pRF model (**A,B**) or a DoG center-surround pRF (**C,D**); however, the ground truth pRF in all these analyses was always a 2D Gaussian pRF. **E-F**. Goodness-of-fit between the estimated pRF shape and the true pRF, separately for different true pRF sizes (rows) and the three simulated datasets (columns). The left three columns show Canonical fit results and the right three columns show the Concurrent fit results. Results are split between the 2D Gaussian pRF model (**E**) and the DoG center-surround pRF model (**F**).

For pRFs whose ground truth shape was a 2D Gaussian, we found that irrespective of the dataset or analysis, the position (x_0_ and y_0_) estimates were closely aligned to the ground truth, with some small exceptions (which occurred especially for pRFs near the stimulus edge). Also, pRF size estimates were generally close to the ground truth. However, in the Canonical fit, for data synthesized with Vista or the de Haas HRF the analysis tended to underestimate pRF size. In contrast, in the Concurrent fit, these errors were substantially reduced, and the analysis typically closely matched the true pRF size. The only exception occurred for the smallest true pRFs (σ=0.25°), where both fits generally overestimated pRF size.

Because our pRF models are circularly symmetric, we can visualize these estimates as one-dimensional profiles through the pRF center. We selected profiles fit to each simulated iteration of a given ground truth pRF, plotted these profiles relative to the true pRF center position, and then averaged these profiles. For the canonical fit, this revealed considerable misestimates of pRF size especially for small and large true pRFs (Figure 3A). These errors were substantially reduced in the Concurrent fit (Figure 3B). The goodness-of-fit between the pRF estimates and the ground truth is shown in Figure 3E. This demonstrates that the Concurrent fit improved pRF estimates especially for pRF sizes between 0.5° and 2°. We observed qualitatively similar results for data synthesized using different stimulus designs (Supplementary Figure S5). However, data synthesized using the de Haas HRF with a combined wedge-and-ring stimulus produced poorer pRF fits in the Concurrent than the Canonical fit (Supplementary Figure S5A).

### Improved fit for center-surround model

We next tested what happens when fitting a DoG center-surround pRF model when the ground truth pRFs are in fact Gaussian. In addition to pRF position (x_0_ and y_0_), the DoG model is defined by an excitatory center size (σ_1_), an inhibitory surround (σ_2_) size, and a parameter (δ) for the ratio of the amplitudes of the two Gaussians. In the Canonical fit, this resulted in egregious errors: while pRF positions were again estimated relatively accurately, the analysis estimated large and pronounced inhibitory surrounds (Figure 3C). The errors were most pronounced for data synthesized using the de Haas HRF and were generally worse for larger pRFs (Figure 3F). In contrast, in the Concurrent fit the inhibitory surround was consistently estimated as shallower, reflecting the absence of any true inhibitory surround in the pRF profile (Figure 3D). Fits were qualitatively similar for all datasets irrespective of the underlying HRF used to synthesize data, although the model failed to capture the smallest or largest pRFs properly (Figure 3E).

For completeness, we repeated these analyses for ground truth data synthesized with a DoG profile. We only report results for fitting a DoG pRF model, because fitting a Gaussian to such data inevitably results in poor fits. First, we confirmed that this analysis could also yield accurate estimates of the HRF used to synthesize data (Supplementary Figures S6-S8). We then again compared pRF estimates against the ground truth. In the Canonical fit there were again considerable errors, most pronounced for pRFs with large inhibitory surrounds. Generally, the model often misestimated the surround (Supplementary Figure S9). In the Concurrent fit, these errors were considerably reduced (Supplementary Figure S10), at least for relatively shallow or smaller surrounds (δ<0.5). Nevertheless, there were many combinations of pRF parameters where the modelling generally produced poor results, for instance when the inhibitory surround was more than twice as large as the facilitatory center pRF. These findings are summarized in Supplementary Figure S11.

### Empirical HRF fits do not match canonicals

Having established that the Concurrent fit can effectively estimate the ground truth HRF used to simulate data, we next reanalyzed several existing pRF mapping datasets. All these datasets used a TR of 1 s. Specifically, these data were from the HCP 7 Tesla dataset (n=32, 1.6 mm isotropic voxels, sweeping bar stimulus) (Benson et al., 2018; Van Essen et al., 2013), our own Auckland 3 Tesla dataset (n=23, 2.3 mm isotropic voxels, sweeping bar stimulus) (Tangtartharakul et al., 2023), and our London 1.5 Tesla dataset (n=17, 2.3 mm isotropic voxels, combined wedge-and-ring stimulus) (van Dijk et al., 2016). All these analyses fit a classic two-dimensional Gaussian pRF model and we only analyzed data points (vertices in cortical surface mesh) whose goodness-of-fit exceeded R^2^>0.2 in the Canonical fit. Figure 4 shows retinotopic maps from an example participant from the Auckland dataset. While the functional architecture of polar angle and eccentricity was similar in both analyses, pRF sizes were subtly larger in the Concurrent fit.

**Figure 4.**
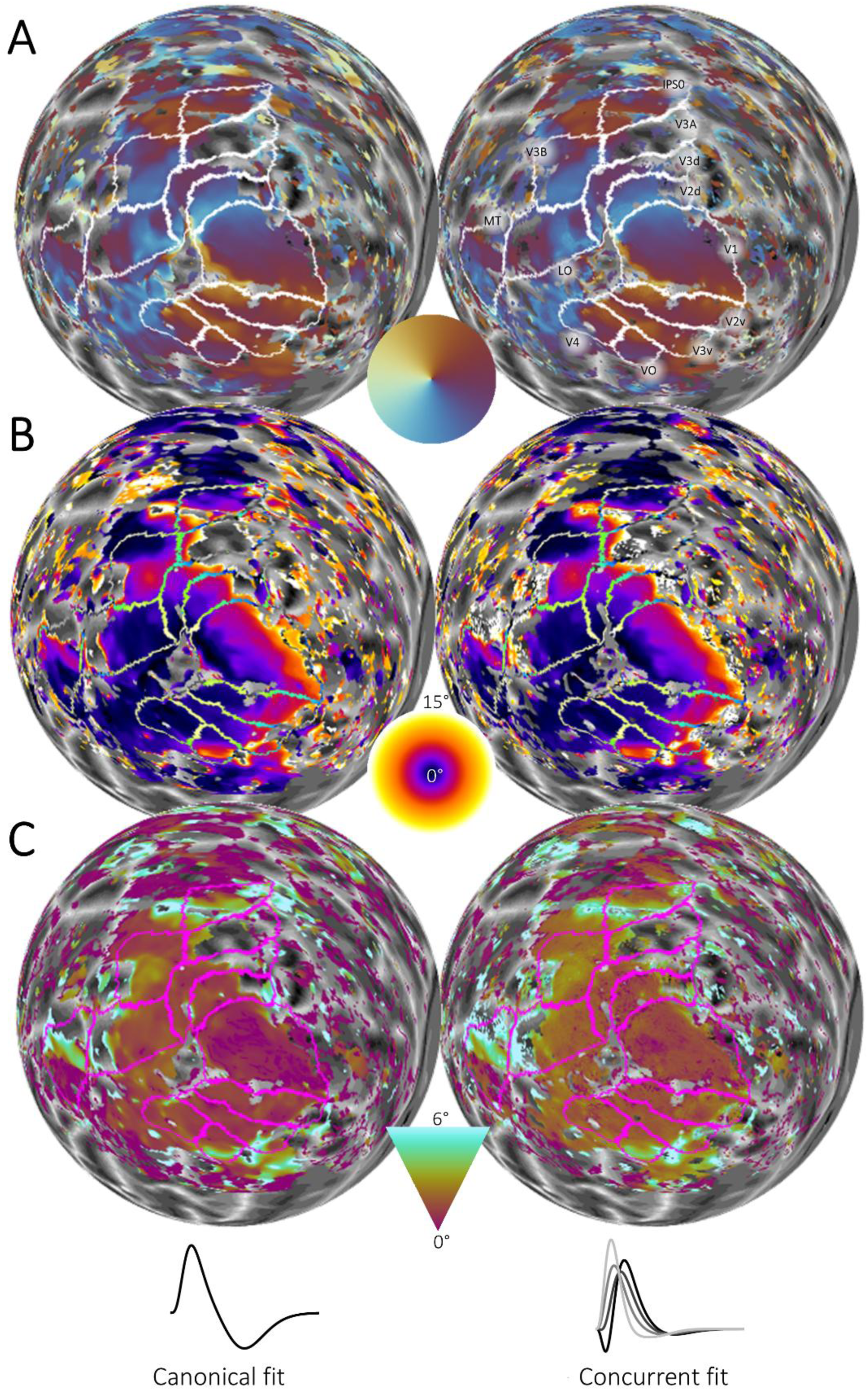
pRF maps of an example participant in the Auckland dataset. Maps of polar angle (**A**), eccentricity (**B**), and pRF size (**C**) estimates are shown on a spherical model of the left cortical hemisphere. Left: maps obtained using Canonical fit. Right: maps obtained using Concurrent fit.

To quantify these results, we averaged the HRF fits separately for each visual region, for all data above threshold. This demonstrated two main points. First, the HRF fit for none of these datasets matched any of our canonical HRFs (Figure 5A-C). Instead, the HRFs estimated from these data tended to peak earlier. They also peaked earlier than the impulse-response functions we had estimated empirically (Figure 1I) for the London dataset (van Dijk et al., 2016). Second, the HRFs fit to these three datasets also differed from one another.

**Figure 5.**
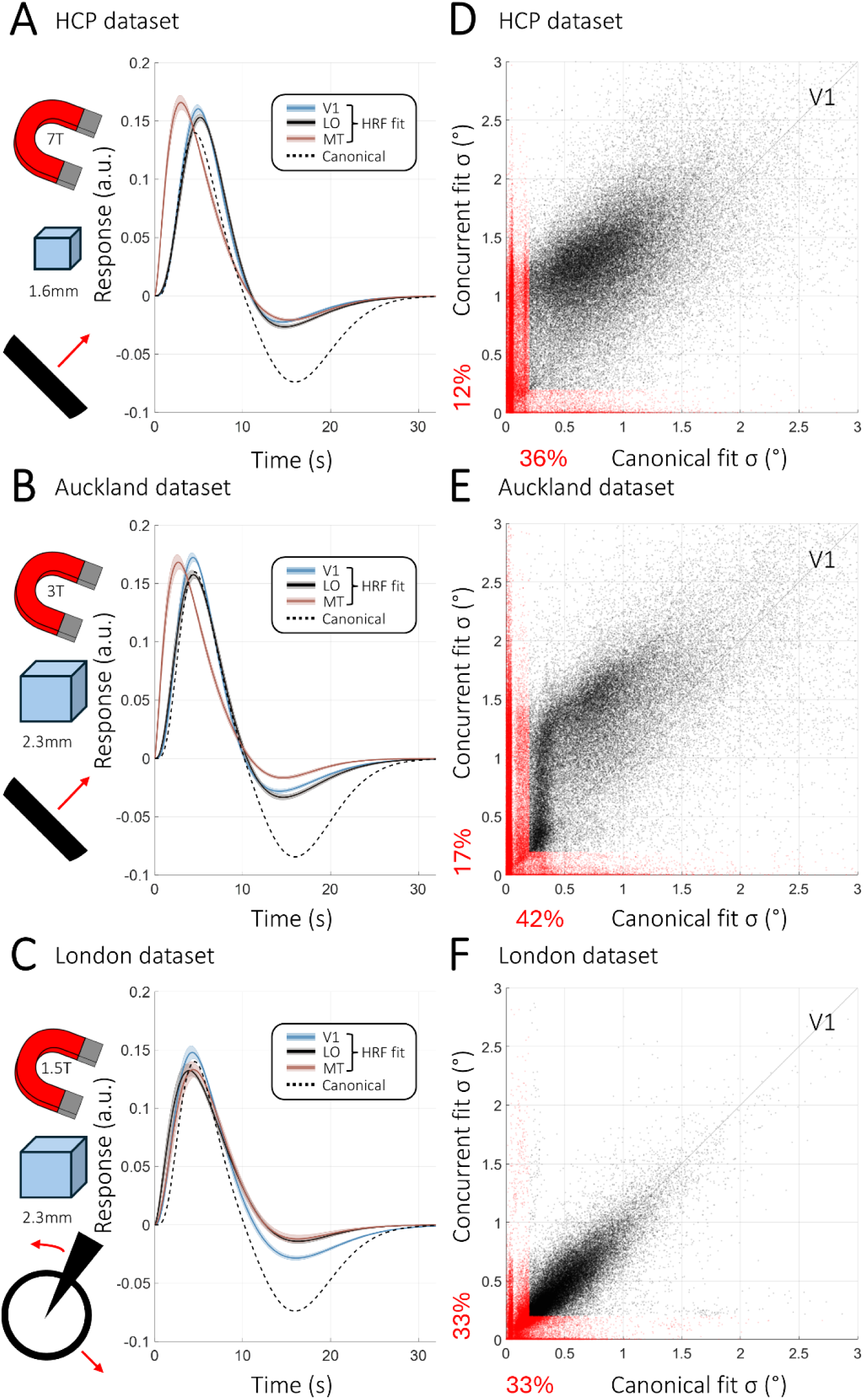
Concurrent HRF fitting of empirical pRF data. **A-C**. Average HRFs estimated in Concurrent fit for the HCP (**A**), Auckland (**B**), and London (**C**) datasets. HRFs were averaged separately for areas V1, LO, and MT, and then averaged across participants (shaded regions denote ±1 standard error of the mean across participants). The dashed black line denotes the SamSrf de Haas canonical HRF for comparison. **D-F**. pRF size estimates for V1 from the Concurrent fit plotted against the corresponding estimates from the Canonical fit (estimates with R^2^>0.2 only). Data are shown for the HCP (**D**), Auckland (**E**), and London (**F**) datasets. Data were pooled across participants in each dataset. The data points labelled in red are those with implausibly small pRFs (σ<0.2°) in an analysis. The red numbers denote the percentage of these pRFs.

### HRF estimates differ between visual regions

Specifically, when we averaged HRF fits separately for each delineated region of interest we revealed that for the two datasets acquired using sweeping bar stimuli there were differences in HRFs estimated across visual cortex (Figure 5A,B). HRF fits in early visual cortex (such as V1-V3) peaked significantly later than those in most of the higher extrastriate regions, like V3A and MT (repeated measures analysis of variance for HCP dataset: F(9,279)=116.57, p<0.0001; Auckland: F(9,198)=84.86, p<0.0001). For the London data, acquired using a combined wedge-and-ring stimulus, there were also significant regional differences (F(9,144)=19.45, p<0.0001). However, the pattern of results was not the same: for example, HRFs in MT and V1 peaked at similar latencies (paired t-test: t(16)=-0.79, p=0.4406), although the lack of significance does not rule out more subtle differences. In LO, the HRF peaked slightly earlier than in V1 (t(16)=2.53, p=0.0224), and ventral regions V4 and VO peaked significantly earlier than either V1 and MT (all p<0.0001). The undershoot was most pronounced in V1 compared to the other regions for the Auckland and London datasets, while such differences were negligible in the HCP dataset (Figure 5A-C). HRF fits to all visual regions we analyzed are shown in Supplementary Figure S12.

### Cross-validation analysis

Unlike in our simulation analyses, for empirical data the ground truth is unknown. In noisy data from real participants there is a risk of overfitting when applying a more complex model like the Concurrent HRF fit. One way to assess this is by cross-validation on split half data. For this purpose, we focused on the Auckland 3T dataset because this showed the most pronounced differences between the HRF fits for different visual regions. We split six runs acquired from the same session into odd- and even-numbered runs, averaged the two sets separately, and then analyzed each of these cross-validation folds with both the Concurrent and the Canonical fit. All other analysis parameters were identical as for the full Auckland 3T dataset.

To determine how consistent the concurrent fit estimated the underlying HRF in the data, we averaged the HRF fits for different visual areas separately for the two independent cross-validation folds. The resulting HRF fits were a close match between folds, again revealing a distinct difference in the peak latency for MT compared to V1 or LO (Supplementary Figure S13).

Next, we determined whether Concurrent fit is superior to the Canonical fit in terms of predicting the other fold’s observed time series. We calculated the cross-validated goodness-of-fit (cR^2^) for each analysis by the fit between the predicted time series from one dataset against the empirically observed time series from the independent, left-out runs. This revealed that the Concurrent fit consistently produced higher cR^2^ values than the Canonical fit. Comparing this for each of the 10 brain regions and 23 individual participants revealed significantly improved fits (Wilcoxon signed rank test: p<0.001, Bonferroni corrected) in all tests, except for three participants in MT and one participant in IPS0. This improvement was especially pronounced for vertices where the overall cR^2^ was greater than zero (Supplementary Figure S14). Negative cR^2^ arise with very noisy data where the fit is poor. We compared participant-wise median cR^2^ across the whole analysed cortex after excluding vertices with negative mean cR^2^. This again revealed consistently better model fits for the Concurrent fit in all but one participant, a highly significant difference (grand mean for Canonical fit M_can_=0.11; Concurrent fit M_con_=0.14; paired t-test: t(22)=7.43, p<0.0001). This indicates that when there is a sufficient signal to be estimated, the Concurrent fit consistently outperformed the Canonical fit.

### Fewer implausibly tiny pRFs

We also compared results for the three empirical datasets (HCP, Auckland, and London datasets) for the Concurrent and Canonical fits. Plotting the estimated pRF size for the fits against one another revealed that at least in the Auckland and the HCP datasets, we observed a considerably smaller proportion of tiny pRFs (σ<0.2°) in the Concurrent than the Canonical fits (Figure 5D,E). Such small pRFs are considerably smaller than the width of the bar stimulus used for mapping (∼0.5° in both datasets), resulting in an under-constrained model. They mostly occur for the early visual regions (Figure 6A) where pRF sizes are smaller, such as V1 (Dumoulin & Wandell, 2008). In contrast, in the London dataset the proportion of such small pRFs was similar in both fitting analyses (Figure 6A).

**Figure 6.**
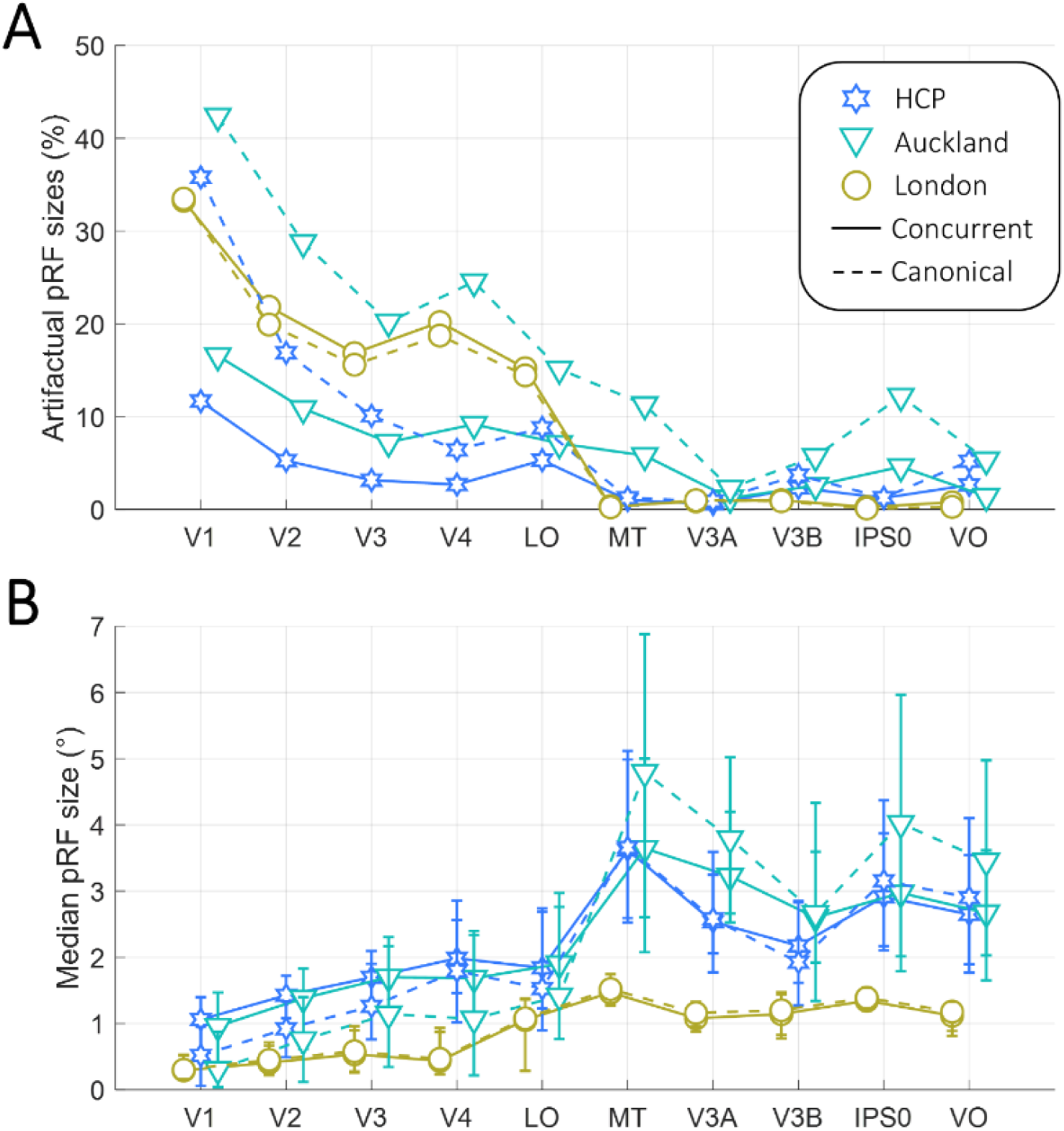
pRF size estimates. The percentage of pRFs with implausibly small pRFs (σ<0.2°) in an analysis (**A**) and median pRF sizes (**B**) are plotted for each visual region and three empirical datasets. Solid lines: Concurrent fit. Dashed lines: Canonical fit. Error bars in B denote the interquartile range of pRF size estimates pooled across participants.

In fact, in both datasets collected using sweeping bar stimuli, pRF sizes tended to be generally larger in the Concurrent than the Canonical fit (Figure 5D,E; Figure 6B). In contrast, in the London dataset collected using a combined wedge-and-ring stimulus, pRF sizes were generally similar for the two fitting analyses. However, they were also substantially smaller than those obtained in the other two datasets (Figure 5F; Figure 6B).

### Comparing pRFs between experiment conditions

Finally, we revisited another empirical data set (Altan, Morgan, et al., 2025). In that study, we reported that adaptation to high spatial frequency patterns yields larger pRF size estimates than adapting to low spatial frequency patterns. We interpreted this as evidence that pRF size is linked to spatial frequency tuning of neuronal populations. However, that analysis assumed the canonical de Haas HRF. This left the possibility that the HRF changes under the different experimental conditions, confounding that result.

We now reanalyzed this dataset with the Concurrent HRF fit. The overall result is qualitatively the same: across all visual regions tested, adapting to high spatial frequency produces larger pRFs than adapting to low spatial frequency (Figure 7A). Numerically, the estimates show far less inter-individual variability, especially in the higher visual regions. On the other hand, pRFs in the early visual cortex (V1-V3) tended to be larger than in the Canonical fit; this is probably partly because the Concurrent fit produced fewer implausibly tiny pRF estimates, just as in our analyses of the Auckland 3T and HCP 7T data. Moreover, one participant who, in the original analysis, bucked the group trend in V1 and had larger pRFs after low-spatial frequency adaptation in the Canonical fit, showed results consistent with the rest of the group in the Concurrent fit.

**Figure 7.**
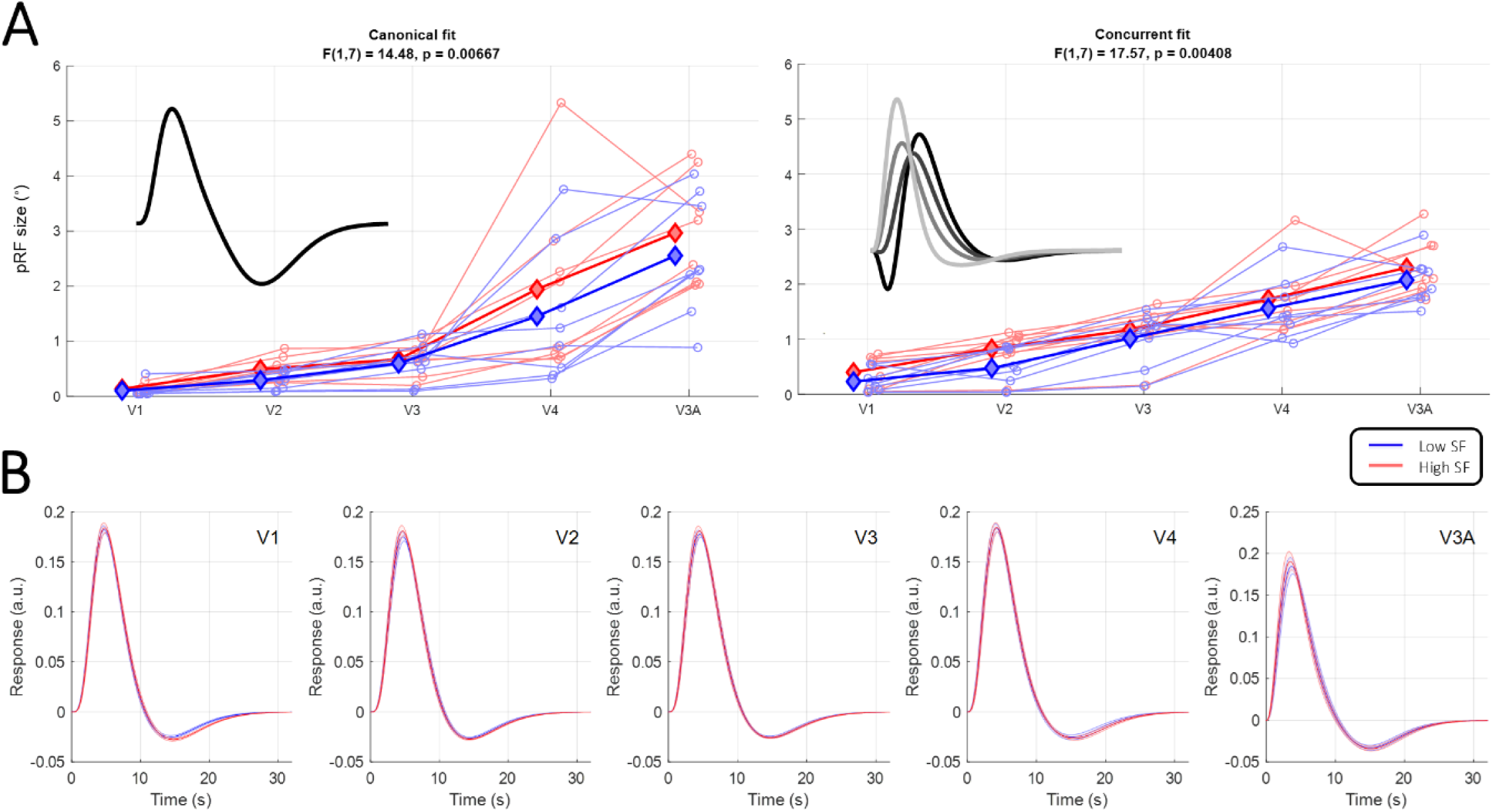
Reanalysis of experimental differences in pRF size. **A**. *Left:* Data from a previous study (Altan, Morgan, et al., 2025) showing adaptation to high-(red) vs. low-spatial frequency (blue) patterns, respectively, yields larger vs. smaller pRF size estimates across human visual cortex. These data were originally analyzed using the Canonical fit. *Right:* The same data but reanalyzed using Concurrent HRF fit. Thin lines and circles denote median pRF size for individual participants. Thick lines and diamonds denote the group mean. Statistics at the top show main effect of spatial frequency adaptation on a two-way repeated measures analysis of variance. **B**. HRF estimates from this Concurrent fit plotted for each visual region separately for high (red) and low (blue) spatial frequency adaptation conditions. Solid line shows the mean HRF across participants. The shaded regions denote ±1 standard error of the mean across participants.

Importantly, the HRF estimated for the two conditions were similar (Figure 7B). Even if there may be subtle differences between the conditions in different visual regions, these are inconsistent between regions and are generally an order of magnitude smaller than the HRF variability we observed between visual regions or between datasets.

## DISCUSSION

We demonstrate that it is possible to estimate the HRF concurrently as part of fitting pRF models. For maximal flexibility, we used a complex model with five free parameters to describe the HRF. As even the simplest pRF models have three parameters this entailed estimating a multidimensional parameter space. We leveraged the conventional coarse-to-fine fitting approach commonly used in pRF analysis (Dumoulin & Wandell, 2008): we first use an extensive grid search of the simple pRF model assuming a canonical HRF from SPM (Friston et al., 1998), and then use parameter optimization to home in on the best parameters for both the pRF and the HRF.

Using simulated data, we found that this approach closely estimated the HRF used to synthesize data for a range of stimulus designs – provided the TR of the simulated data was 1 s. For longer TRs, the HRF fit became considerably worse, possibly because estimating the HRF shape accurately requires good temporal resolution. In this context, we note that empirically estimated impulse-response functions showed marked differences between different visual regions (Figure 1I) when measured with a TR of 1 s (van Dijk et al., 2016), but we previously had not observed such differences with longer TRs (Schwarzkopf et al., 2014). With the advent of accelerated imaging (Breuer et al., 2005), short TRs have become commonplace in the pRF literature (Benson et al., 2018; Chang et al., 2025; Harvey & Dumoulin, 2011; Himmelberg et al., 2022; Morgan & Schwarzkopf, 2019; Moutsiana et al., 2016; Tangtartharakul et al., 2023; van Dijk et al., 2016). In situations necessitating longer TRs, it could be possible to model the time series with a finer temporal resolution than the scan TR. This would entail using a smoothly changing stimulus design where stimulus onset is not time-locked with image acquisition (Benson et al., 2018; Chang et al., 2025). Such efforts are beyond the scope of the present study; our further analyses focused only on designs with TR of 1 s.

To test the relevance of modelling the HRF in real data, we chose four empirical datasets, three from our own studies (Altan, Morgan, et al., 2025; Tangtartharakul et al., 2023; van Dijk et al., 2016) and a subset of data from the HCP (Benson et al., 2018), to compare pRF results with and without concurrent HRF fitting. Our pRF estimates were similar between canonical and concurrent HRF fitting. But critically, these analyses also suggested important differences that could affect the interpretation of pRF studies. Importantly, we found that the best-fitting HRF for these empirical data differed from all the canonical HRF functions and from individual HRFs estimated empirically as impulse-response functions. They especially differed in terms of response latency. Assuming a fixed canonical HRF could therefore have significant consequences on pRF parameter estimates in published studies.

We concede that in empirical data without a known ground truth, we cannot tell if (and how) model parameters trade off against one another. Especially in classical mapping paradigms, the temporal parameters of the HRF are confounded with the temporal sequence of the stimulation protocol. As such, spatial parameters like pRF size could trade off against temporal parameters of the HRF, such as its latency and dispersion (Dumoulin & Wandell, 2008). Along a similar vein, the depth of the HRF undershoot could confound estimates of inhibitory interactions in complex pRF models (Anderson et al., 2017; Aqil et al., 2021, 2025; Harvey et al., 2013; Schwarzkopf et al., 2014; Zuiderbaan et al., 2012). Our cross-validation analysis of empirical data suggests model fits improved under the concurrent HRF fit. This suggests there was minimal overfitting when including the five additional free parameters of the HRF, although this cannot categorically rule out issues with identifying parameters. However, the accuracy of the concurrent fit in our simulations demonstrated that at least in synthetic data, using a 2D Gaussian pRF model and a TR or 1 s, pRF and HRF parameters could be estimated with high precision, suggesting that this problem is not severe under these conditions and variability of HRF and pRF parameters can be reliably distinguished. Spot tests of randomly selected vertices suggest that the Concurrent fit generally arrives at the optimal combination of parameters to fit to time series. We acknowledge that testing this systematically would require an extensive analysis that is outside the scope of the present study.

### Substantial effects of HRF on pRF estimates from bar stimuli

Our simulation analyses revealed that failing to account for the correct underlying HRF profoundly impacts pRF parameter estimates. Specifically, when we assumed the fixed canonical HRF we had used in previous studies (Anderson et al., 2017; de Haas et al., 2014; Morgan & Schwarzkopf, 2019; Moutsiana et al., 2016, 2016; Schwarzkopf et al., 2014; Tangtartharakul et al., 2023), small pRF sizes – as are typically found in early visual cortex – were generally underestimated. This accords with previous observations of incorrect pRF size estimates when assuming the wrong HRF (Lage-Castellanos et al., 2020; Lerma-Usabiaga et al., 2020). It is also consistent with our findings from empirical data: in early visual regions up to V4, concurrent HRF fitting consistently yielded larger pRFs than assuming a fixed HRF, at least when using a sweeping bar stimulus.

Crucially, this general trend also meant we observed substantially fewer implausibly tiny pRFs, in particular in the earlier visual regions where pRFs tend to be smaller (Dumoulin & Wandell, 2008). Our definition of what constitutes artifactual small pRFs (σ<0.2°) is arbitrary, but such pRFs would have been considerably smaller than the width of the bar stimulus. This means their position is ambiguous and cannot be accurately captured by the standard pRF analysis. Again, one potential solution for estimating such tiny pRFs – if they exist – is to use a smoothly drifting bar stimulus and model the time series at a finer temporal resolution than the TR. However, many of these tiny pRFs were considerably smaller than our criterion, often considerably smaller even than 0.1°. It seems biologically implausible that there should be many such pRFs outside the foveal confluence. Our present results suggest that by capturing the shape of the HRF in the model the analysis can help disambiguate the responses of such voxels more accurately than when assuming a fixed HRF.

### Minimal effects of HRF for combined wedge-and-ring stimuli

With the combined wedge-and-ring stimulus we had used in previous studies (Alvarez et al., 2015; Anderson et al., 2024; Farahbakhsh et al., 2022; Moutsiana et al., 2016; van Dijk et al., 2016), we also found that concurrently modelling the HRF somewhat improved pRF parameter estimates (i.e., relative to ground truth). However, we found no difference in pRF sizes between the two analyses in our empirical wedge-and-ring dataset. Importantly though, pRFs in this dataset were estimated as considerably smaller than those obtained for the other two datasets using bar stimuli.

In an earlier study, we had found the combined wedge-and-ring design was more efficient than sweeping bar stimuli, due to better duty cycles and more estimates passing the variance explained threshold, and that it tended to estimate smaller pRF sizes (Alvarez et al., 2015). Our present finding of smaller pRFs with the wedge-and-ring stimuli confirms this and also agrees with previous findings that using polar wedge and eccentricity ring stimuli affects pRF size estimates (Linhardt et al., 2021) (but note that this earlier work did not combine the stimuli in the same run as we did). Specifically, that study found that relative to using sweeping bar stimuli, pRFs near the central visual field were estimated more reliably and with smaller sizes. This likely reflects the fact that polar wedges can estimate small pRFs more accurately, especially when the pRF is substantially smaller than the bar width and the bar steps through the visual field time-locked with image acquisition (but again see our suggestion of addressing this problem using methods with a finer temporal resolution). However, for combined wedge-and-ring stimuli we also observed somewhat greater errors of pRF estimates relative to ground truth parameters in our simulation analyses. Taken together, these differences suggest caution when using wedge-and-ring stimuli for pRF estimation. Interestingly, recent work proposed a novel warped bar stimulus whose width scales logarithmically with eccentricity; this improves the reliability of pRF size estimates (Chang et al., 2025) and may offer a good compromise for estimating pRFs reliably across the visual field.

### Dramatic consequences of HRF for estimating complex pRF models

Our simulation analyses also showed that failing to account for differences in the HRF can dramatically impact the estimates for more complex pRF models. While most pRF studies still use the conventional two-dimensional Gaussian model, several more refined models have been employed. Approaches range from inhibitory center-surround interactions (Zuiderbaan et al., 2012), elliptic pRFs (Lerma-Usabiaga et al., 2021; Silson et al., 2018), modelling nonlinear spatial summation of the fMRI response (Kay et al., 2015; Kay, Winawer, Mezer, et al., 2013; Kay, Winawer, Rokem, et al., 2013), and even complex divisive normalization models that encompass several of these processes under the same formulation (Aqil et al., 2021, 2024, 2025). All these models considerably increase the number of parameters the model must estimate, which increases the risk of local minima and spurious findings.

Here, we tested the effect of concurrently modelling the HRF when estimating the DoG model where a larger inhibitory pRF surrounds a central facilitatory component (Zuiderbaan et al., 2012). Such models have previously suggested a link between alpha wave activity and the strength of pRF inhibition (Harvey et al., 2013). We also previously suggested that the inhibitory surround is dampened in schizophrenia (Anderson et al., 2017), but not in autism spectrum disorders (Schwarzkopf et al., 2014). Similar findings (albeit using the even more complex divisive normalization model (Aqil et al., 2021)) have suggested that inhibitory pRF modulation relates to neurotransmitter receptors (Aqil et al., 2024) and the perceptual effects of psychedelics (Aqil et al., 2025). These reports underline the importance of assessing the validity of these complex models.

We found that failing to account for the true HRF used to synthesize data caused profound errors when estimating the complex DoG model. Even though the true pRF was a two-dimensional Gaussian, the analysis suggested strong and widespread inhibitory surround interactions. The canonical de Haas HRF we used has a large undershoot, which may have rendered these errors particularly severe, but this alone does not explain these errors. There was no consistent pattern of errors depending on the true HRF used to synthesize data.

However, when we concurrently modelled the HRF in this analysis the errors were reduced considerably. While the analysis still struggled to estimate the smallest and largest pRFs accurately, the errors were much smaller. Importantly, for most pRFs the model estimated the inhibitory surround as very shallow, reflecting the fact that there was no inhibitory surround in the true pRF. Optimally, one would expect that the model estimates the inhibitory pRF component as having zero amplitude – however, as amplitude is zero bounded, it is unsurprising that the average amplitude across multiple noisy data points will be greater than zero.

We also tested the performance of the analysis when the true pRFs had a center-surround profile. Here, again, we found that concurrently modelling the HRF improved parameter estimates compared to assuming a fixed canonical HRF. Importantly, however, many significant errors remained in these analyses. This spells further caution for the use of such complex models. We note though that these analyses may represent a worst-case scenario. We estimated our DoG model using the coarse-to-fine fitting approach where we first approximate the five parameters of the DoG pRF via a grid search algorithm. We did this to match conditions between the DoG and conventional 2D Gaussian analyses, but due to computational constraints this necessitated using a relatively sparse search grid. This approach differs from how we have previously used the DoG model (Anderson et al., 2017; Schwarzkopf et al., 2014) or how others have estimated complex pRFs (Aqil et al., 2021): typically those studies start with a simpler pRF model to approximate initial parameters about pRF position and size with high accuracy, and then extend the model from those estimates. One potential approach for dealing with this could be to use a standard 2D Gaussian model to first concurrently model the HRF at each voxel or vertex in the cortical surface mesh and then use these HRF fits when estimating the complex pRF models. Such an iterative approach may improve the accuracy of complex model fits, but it could also run the risk of local minima. It would be valuable to quantify this, but this is beyond the scope of our present study. Generally, our findings here emphasize that a valid interpretation of such complex models should always go alongside ground truth simulations of the analysis approach used (Binda et al., 2013; Chang et al., 2025; Lerma-Usabiaga et al., 2020; Senden et al., 2014; Zeidman et al., 2017).

In this context we reiterate that our present analysis also includes an iterative approach: during the grid search stage of the analysis, we assumed the SPM canonical HRF as perhaps the “most canonical” example (note that the Vista HRF is quite similar to the SPM function). We then used its default parameters as starting points in the optimization stage. This did not affect the HRF fits dramatically in most analyses – but it is probably why HRF fits tended to be most accurate when the true HRF used for synthesizing data was based on the SPM HRF. Future research should quantify the impact of the initial HRF parameters. It constitutes a large-scale endeavor because it requires a substantial number of permutations of analyses.

### HRF estimates differ between visual regions and stimulus designs

We also observed pronounced differences in HRF shape across the different visual areas, although this also depended on the dataset: in the HCP and Auckland datasets, the HRF peaked considerably earlier in the MT complex and higher dorsal regions like V3A and IPS0 than in V1-V3. At first glance, this finding may seem counterintuitive: considering that these regions are later in the visual processing hierarchy, one would expect the response to peak later than in earlier regions like V1. However, the difference in neuronal response latencies should be on the order of a few hundred milliseconds, considerably less than the 1-2 s differences we observed. As such, neurovascular factors or spatiotemporal response dynamics are a better explanation for our findings. In fact, even pRF size could explain such differences: because higher regions usually have much larger pRFs, a voxel might be distally activated by the stimulus and so the signal will not completely decay back to baseline (except for prolonged blank screen epochs). This continuous activation could manifest as earlier peaks in the concurrently estimated HRF. However, our simulations suggest that this is unlikely to be the case: synthetic data from large pRFs do not show any systematic differences in the peak latency estimates compared to small pRFs (Supplementary Figures S3 and S4).

In this context, it is interesting to note that previous reports of regional variation in HRFs also suggest that higher-level regions like the frontal and supplementary eye fields may peak earlier than primary visual and motor cortex (Handwerker et al., 2004). However, these differences are highly idiosyncratic and are from anatomically far more distinct brain areas than the visual regions we investigated across occipitotemporal cortex.

In any case, the variability in HRFs we observe probably does not only reflect variability in the neurovascular HRF but are a description of fMRI responses within a given brain region in the experimental context of a pRF experiment. In this light it is interesting that HRFs in the London dataset in early visual cortex and MT and other dorsal regions were not significantly different, but category-selective regions in lateral occipital and ventral occipital cortex peaked earlier instead. These three datasets differed in several respects, including the stimulus design and materials, magnetic field strength, and voxel size. It is therefore difficult to determine what caused these differences in the HRF.

However, the most parsimonious explanation may be that the HCP and Auckland data were acquired using sweeping bar stimuli while the London dataset used a combined wedge-and-ring stimulus. While differences in magnetic field strength could affect the contribution of macro-vs. micro-vasculature in different datasets, it is not obvious why this should result in regional differences in response latencies. Moreover, in the London dataset we had also collected impulse-response functions using full-screen stimuli in a slow event-related design. This revealed marked regional differences of around 1.5 s between MT and early visual cortex akin to what we observed for the HCP and Auckland data (Figure 1I). The fact we did not observe the same pattern of results for the concurrent HRF fits in the London data therefore suggests that estimated HRF shape also depends on the specific pRF stimulus paradigm.

Indeed, this is unsurprising because hemodynamic responses are non-linear (Lindquist et al., 2009). Most pRF mapping studies use paradigms in which a stimulus traverses the visual field in an orderly and predictable sequence. With a sweeping bar stimulus, a participant can have clear expectations as to when the bar will reach a given location. However, in previous work we showed that predictability does not majorly impact pRF estimates in visual cortex – rather, differences between stimulus sequences are predominantly due to their spatiotemporal characteristics (Infanti & Schwarzkopf, 2020). Ordered designs must induce spatiotemporal correlations across the cortical surface. These issues could theoretically be addressed by using unpredictable, random stimulus sequences. However, random sequences typically produce lower signal-to-noise ratios (Binda et al., 2013), possibly because they overlap more with thermal and other noise sources in the fMRI signal.

Non-linear summation of the signal (Kay, Winawer, Mezer, et al., 2013; Kay, Winawer, Rokem, et al., 2013) will also affect responses especially in the combined wedge-and-ring stimulus design. Unlike with the sweeping bar stimuli there is continuous stimulation across distal parts of the visual field in this design. Nevertheless, these points remain speculative; to understand the HRF differences between our three datasets will require a more systematic comparison. Compressive summation is also more pronounced in higher extrastriate regions and could thus account for neural responses peaking earlier there. Others have modeled such nonlinearities directly in the pRF model (Kay, Winawer, Mezer, et al., 2013), but this involves applying an exponent to the pRF response output, which precludes modelling inhibitory interactions like in the DoG model. While the even more complex divisive normalization model can capture both compressive nonlinearity and inhibitory interactions (Aqil et al., 2021), it requires a substantial reorganization of pRF modelling process by combining two pRFs, plus additional scaling factors.

In contrast, if these interactions can instead be captured by allowing flexibility in the HRF this considerably simplifies the analysis. This can be desirable, at least in studies that seek to estimate spatial selectivity of voxels (pRF size, plus potentially inhibitory surround interactions). However, this could come at the expense of information that these complex models might potentially provide (Aqil et al., 2024). Future investigations should conduct direct comparisons to benchmark these different approaches to understand the trade-offs between these models – and if concurrent HRF estimation can still help in these scenarios. In any case, in the context of standard pRF models assuming a constant impulse-response function may be suboptimal, regardless of whether this is a canonical function or one estimated for each individual participant.

### Reanalyzing experimental conditions

The HRF might also vary between different experimental conditions or task demands. We recently published a pRF study using our standard analysis assuming a canonical HRF (Altan, Morgan, et al., 2025). Reanalyzing these data with the Concurrent fit replicated our original finding of larger pRFs after adapting to high-vs low-spatial frequency patterns. Notably, pRF size estimates however showed less inter-individual variability and there were fewer implausibly small pRFs. Importantly, the HRFs estimated for the two experimental conditions were virtually identical. Naturally, as these are empirical data, the ground truth is unknown. However, combined with our simulations, our empirical results, and the cross-validation results, this finding implies that the adaptation condition did not modulate the HRF; rather, accounting for the HRF shape in the model presumably reduced noisy estimates.

### Other ways to estimate HRF

To our knowledge, our study is the first to describe the variation of HRF shapes in pRF mapping paradigms. However, other recent studies also modelled the HRF concurrently with the pRF model for each voxel or brain coordinate. Specifically, one study proposed a Bayesian algorithm for pRF estimation (Zeidman et al., 2017). This method leveraged the algorithmic engine underlying dynamic causal modelling (Friston et al., 2003), including the balloon model of hemodynamic responses (Buxton et al., 2004; Stephan et al., 2007).

We note that all our analyses are based on a shared family of HRF, the widely used double-gamma function as HRF (Boynton et al., 1996; Friston et al., 1998). It is possible that this constrained our findings to similar HRF shapes. It may therefore fail to account for very different underlying basis functions, a limitation that could be particularly problematic outside the cortex. On the other hand, by fitting the HRF with five free parameters we allowed substantial flexibility in the shape; for example, the HRF could theoretically capture an initial dip in the function through a broad undershoot. Nevertheless, future research could readily extend this framework to other HRF models.

Other recent studies have modelled differences in HRF shape using a spline algorithm (Kay, Winawer, Mezer, et al., 2013) or by fitting temporal derivatives of the canonical HRF (Aqil et al., 2024, 2025). The latter approach reduces the number of free parameters that must be estimated – the differences in response latency that characterize most of the variation we observed in our present work can probably be captured using only *one* additional parameter in the model. Especially when fitting complex pRF models, this could be beneficial. However, to capture differences in dispersion requires a second parameter, while modelling potential variability in undershoot amplitude requires a third. Understanding the potential impact of these assumptions is not trivial. It again underlines the importance of assessing the validity of any approach through ground truth simulation.

It could also be feasible to circumvent the need for explicit HRF fitting, especially in relatively standard analyses on healthy, neurotypical participants. One could instead use a probabilistic atlas of how the HRF varies across the brain and thus constrain the HRF for any given brain coordinate. Such an atlas could be based on group averages, but perhaps a better approach would involve deep learning model predictions. This approach has already been demonstrated to accurately predict idiosyncrasies in retinotopic map architecture (Ribeiro et al., 2021, 2023). The model could be extended to also predict HRF parameters. However, as our present work suggests, this may depend on the stimulus design and scanning parameters.

### Conclusions and recommendations

We showed that differences in the underlying HRF can have dramatic consequences on pRF modelling results. Especially for estimating complex pRF models, assuming a fixed HRF can skew estimates considerably. At least for stimulus designs with short TR, concurrent fitting of the HRF can improve the validity of pRF models. In general, we observed that HRFs estimated by the model can vary between experimental designs and brain regions. The reason for this is probably not merely due to hemodynamics; rather this likely also reflects non-linearity in the response. While such effects could be captured by more complex pRF models, our simulations demonstrated that failure to account for variability in the HRF can dramatically skew complex pRF model estimates.

This underlines the importance of accounting for HRF differences in pRF studies. We recommend using concurrent HRF fitting at least as a robustness check, especially in studies comparing pRFs between different experimental conditions or different groups of participants that might conceivably differ in their HRF or spatiotemporal dynamics of the fMRI response. Moreover, when fitting highly complex pRF models, it serves to conduct simulations with different ground truth HRFs. If this suggests the HRF has a significant impact, it could help to run a concurrent HRF fit via the standard 2D Gaussian model and then incorporate this HRF estimate in the final model.

## ACKNOWLEDGEMENTS

We thank Garikoitz Lerma-Usabiaga, David Linhardt, and Fernanda Ribeiro for constructive discussions of this work.

## SUPPLEMENTARY INFORMATION

Supplementary Figures are available at https://doi.org/10.17605/OSF.IO/9V845.

## DATA, SCRIPT, AND CODE AVAILABILITY

All processed pRF data, raw data time series, and analysis code necessary for reproducing these results are available at https://doi.org/10.17605/OSF.IO/9V845, except for the data reanalyzed from Altan et al. 2025b, which are available at https://doi.org/10.17605/OSF.IO/9KFGX.

## REFERENCES

Aguirre, G. K., Zarahn, E., & D’esposito, M. (1998). The variability of human, BOLD hemodynamic responses. NeuroImage, 8(4), 360–369. 10.1006/nimg.1998.0369

Altan, E., Boyaci, H., Dakin, S. C., & Schwarzkopf, D. S. (2025). Perceiving object size in pictures involves high-level processing. Proceedings of the Royal Society B: Biological Sciences, 292(2046), 20242967. 10.1098/rspb.2024.2967

Altan, E., Morgan, C. A., Dakin, S. C., & Schwarzkopf, D. S. (2025). Spatial frequency adaptation modulates population receptive field sizes. eLife, 13, RP100734. 10.7554/eLife.100734

Alvarez, I., De Haas, B. A., Clark, C. A., Rees, G., & Schwarzkopf, D. S. (2015). Comparing different stimulus configurations for population receptive field mapping in human fMRI. Frontiers in Human Neuroscience, 9, 96. 10.3389/fnhum.2015.00096

Amano, K., Wandell, B. A., & Dumoulin, S. O. (2009). Visual field maps, population receptive field sizes, and visual field coverage in the human MT+ complex. Journal of Neurophysiology, 102(5), 2704–2718. 10.1152/jn.00102.2009

Anderson, E. J., Dekker, T. M., Farahbakhsh, M., Hirji, N., Schwarzkopf, D. S., Michaelides, M., & Rees, G. (2024). fMRI and gene therapy in adults with CNGB3 mutation. Brain Research Bulletin, 215, 111026. 10.1016/j.brainresbull.2024.111026

Anderson, E. J., Tibber, M. S., Schwarzkopf, D. S., Shergill, S. S., Fernandez-Egea, E., Rees, G., & Dakin, S. C. (2017). Visual Population Receptive Fields in People with Schizophrenia Have Reduced Inhibitory Surrounds. Journal of Neuroscience, 37(6), 1546–1556. 10.1523/JNEUROSCI.3620-15.2016

Aqil, M., de Hollander, G., Vreugdenhil, N., Knapen, T., & Dumoulin, S. O. (2025). Psilocybin alters visual contextual computations. Nature Communications, 16(1), 10271. 10.1038/s41467-025-65150-y

Aqil, M., Knapen, T., & Dumoulin, S. O. (2021). Divisive normalization unifies disparate response signatures throughout the human visual hierarchy. Proceedings of the National Academy of Sciences of the United States of America, 118(46), e2108713118. 10.1073/pnas.2108713118

Aqil, M., Knapen, T., & Dumoulin, S. O. (2024). Computational model links normalization to chemoarchitecture in the human visual system. Science Advances, 10(1), eadj6102. 10.1126/sciadv.adj6102

Benson, N. C., Jamison, K. W., Arcaro, M. J., Vu, A. T., Glasser, M. F., Coalson, T. S., Van Essen, D. C., Yacoub, E., Ugurbil, K., Winawer, J., & Kay, K. (2018). The Human Connectome Project 7 Tesla retinotopy dataset: Description and population receptive field analysis. Journal of Vision, 18(13), 23. 10.1167/18.13.23

Binda, P., Thomas, J. M., Boynton, G. M., & Fine, I. (2013). Minimizing biases in estimating the reorganization of human visual areas with BOLD retinotopic mapping. Journal of Vision, 13(7), 13. 10.1167/13.7.13

Boynton, G. M., Engel, S. A., Glover, G. H., & Heeger, D. J. (1996). Linear systems analysis of functional magnetic resonance imaging in human V1. The Journal of Neuroscience: The Official Journal of the Society for Neuroscience, 16(13), 4207–4221.

Breuer, F. A., Blaimer, M., Heidemann, R. M., Mueller, M. F., Griswold, M. A., & Jakob, P. M. (2005). Controlled aliasing in parallel imaging results in higher acceleration (CAIPIRINHA) for multi-slice imaging. Magnetic Resonance in Medicine, 53(3), 684–691. 10.1002/mrm.20401

Buxton, R. B., Uludağ, K., Dubowitz, D. J., & Liu, T. T. (2004). Modeling the hemodynamic response to brain activation. *NeuroImage*, Mathematics in Brain Imaging, 23, S220–S233. 10.1016/j.neuroimage.2004.07.013

Chang, K., Fine, I., & Boynton, G. M. (2025). Improving the reliability and accuracy of population receptive field measures using a logarithmically warped stimulus. Journal of Vision, 25(1), 5. 10.1167/jov.25.1.5

Dale, A. M., Fischl, B., & Sereno, M. I. (1999). Cortical surface-based analysis. I. Segmentation and surface reconstruction. NeuroImage, S(2), 179–194. 10.1006/nimg.1998.0395

de Haas, B., Schwarzkopf, D. S., Anderson, E. J., & Rees, G. (2014). Perceptual load affects spatial tuning of neuronal populations in human early visual cortex. Current Biology: CB, 24(2), R66–67. 10.1016/j.cub.2013.11.061

de Haas, B., Sereno, M. I., & Schwarzkopf, D. S. (2021). Inferior occipital gyrus is organised along common gradients of spatial and face-part selectivity. Journal of Neuroscience. 10.1523/JNEUROSCI.2415-20.2021

Dumoulin, S. O., & Wandell, B. A. (2008). Population receptive field estimates in human visual cortex. NeuroImage, 39(2), 647–660. 10.1016/j.neuroimage.2007.09.034

Engel, S. A., Glover, G. H., & Wandell, B. A. (1997). Retinotopic organization in human visual cortex and the spatial precision of functional MRI. Cerebral Cortex (New York, N.Y.: 1991), 7(2), 181–192.

Farahbakhsh, M., Anderson, E. J., Maimon-Mor, R. O., Rider, A., Greenwood, J. A., Hirji, N., Zaman, S., Jones, P. R., Schwarzkopf, D. S., Rees, G., Michaelides, M., & Dekker, T. M. (2022). A demonstration of cone function plasticity after gene therapy in achromatopsia. Brain, 145(11), 3803–3815. 10.1093/brain/awac226

Fischl, B., Sereno, M. I., & Dale, A. M. (1999). Cortical surface-based analysis. II: Inflation, flattening, and a surface-based coordinate system. NeuroImage, S(2), 195–207. 10.1006/nimg.1998.0396

Friston, K. J., Fletcher, P., Josephs, O., Holmes, A., Rugg, M. D., & Turner, R. (1998). Event-related fMRI: Characterizing differential responses. NeuroImage, 7(1), 30–40. 10.1006/nimg.1997.0306

Friston, K. J., Harrison, L., & Penny, W. (2003). Dynamic causal modelling. NeuroImage, 19(4), 1273–1302.

Handwerker, D. A., Ollinger, J. M., & D’Esposito, M. (2004). Variation of BOLD hemodynamic responses across subjects and brain regions and their effects on statistical analyses. NeuroImage, 21(4), 1639–1651. 10.1016/j.neuroimage.2003.11.029

Harvey, B. M., & Dumoulin, S. O. (2011). The Relationship between Cortical Magnification Factor and Population Receptive Field Size in Human Visual Cortex: Constancies in Cortical Architecture. The Journal of Neuroscience: The Official Journal of the Society for Neuroscience, 31(38), 13604–13612. 10.1523/JNEUROSCI.2572-11.2011

Harvey, B. M., Vansteensel, M. J., Ferrier, C. H., Petridou, N., Zuiderbaan, W., Aarnoutse, E. J., Bleichner, M. G., Dijkerman, H. C., van Zandvoort, M. J. E., Leijten, F. S. S., Ramsey, N. F., & Dumoulin, S. O. (2013). Frequency specific spatial interactions in human electrocorticography: V1 alpha oscillations reflect surround suppression. NeuroImage, 65, 424–432. 10.1016/j.neuroimage.2012.10.020

Himmelberg, M. M., Winawer, J., & Carrasco, M. (2022). Linking individual differences in human primary visual cortex to contrast sensitivity around the visual field. Nature Communications, 13(1), Article 1. 10.1038/s41467-022-31041-9

Infanti, E., & Schwarzkopf, D. S. (2020). Mapping sequences can bias population receptive field estimates. NeuroImage, 211, 116636. 10.1016/j.neuroimage.2020.116636

Kay, K. N., Weiner, K. S., & Grill-Spector, K. (2015). Attention reduces spatial uncertainty in human ventral temporal cortex. Current Biology: CB, 25(5), 595–600. 10.1016/j.cub.2014.12.050

Kay, K. N., Winawer, J., Mezer, A., & Wandell, B. A. (2013). Compressive spatial summation in human visual cortex. Journal of Neurophysiology, 110(2), 481–494. 10.1152/jn.00105.2013

Kay, K. N., Winawer, J., Rokem, A., Mezer, A., & Wandell, B. A. (2013). A two-stage cascade model of BOLD responses in human visual cortex. PLoS Computational Biology, S(5), e1003079. 10.1371/journal.pcbi.1003079

Lagarias, J., Reeds, J., Wright, M., & Wright, P. (1998). Convergence properties of the Nelder—Mead simplex method in low dimensions. SIAM Journal of Optimization, S, 112–147.

Lage-Castellanos, A., Valente, G., Senden, M., & De Martino, F. (2020). Investigating the Reliability of Population Receptive Field Size Estimates Using fMRI. Frontiers in Neuroscience, 14, 825. 10.3389/fnins.2020.00825

Lerma-Usabiaga, G., Benson, N., Winawer, J., & Wandell, B. A. (2020). A validation framework for neuroimaging software: The case of population receptive fields. PLOS Computational Biology, 16(6), e1007924. 10.1371/journal.pcbi.1007924

Lerma-Usabiaga, G., Winawer, J., & Wandell, B. A. (2021). Population Receptive Field Shapes in Early Visual Cortex Are Nearly Circular. The Journal of Neuroscience: The Official Journal of the Society for Neuroscience, 41(11), 2420–2427. 10.1523/JNEUROSCI.3052-20.2021

Lindquist, M. A., Meng Loh, J., Atlas, L. Y., & Wager, T. D. (2009). Modeling the hemodynamic response function in fMRI: Efficiency, bias and mis-modeling. NeuroImage, 45(1 Suppl), S187–198. 10.1016/j.neuroimage.2008.10.065

Linhardt, D., Pawloff, M., Hummer, A., Woletz, M., Tik, M., Ritter, M., Schmidt-Erfurth, U., & Windischberger, C. (2021). Combining stimulus types for improved coverage in population receptive field mapping. NeuroImage, 238, 118240. 10.1016/j.neuroimage.2021.118240

Morgan, C., & Schwarzkopf, D. S. (2019). Comparison of human population receptive field estimates between scanners and the effect of temporal filtering. F1000Research, 8, 1681. 10.12688/f1000research.20496.2

Moutsiana, C., de Haas, B., Papageorgiou, A., van Dijk, J. A., Balraj, A., Greenwood, J. A., & Schwarzkopf, D. S. (2016). Cortical idiosyncrasies predict the perception of object size. Nature Communications, 7, 12110. 10.1038/ncomms12110

Nelder, J. A., & Mead, R. (1965). A Simplex Method for Function Minimization. The Computer Journal, 7(4), 308–313. 10.1093/comjnl/7.4.308

Ribeiro, F. L., Bollmann, S., & Puckett, A. M. (2021). Predicting the retinotopic organization of human visual cortex from anatomy using geometric deep learning. NeuroImage, 244, 118624. 10.1016/j.neuroimage.2021.118624

Ribeiro, F. L., York, A., Zavitz, E., Bollmann, S., Rosa, M. G. P., & Puckett, A. (2023). Variability of visual field maps in human early extrastriate cortex challenges the canonical model of organization of V2 and V3. eLife, 12, e86439. 10.7554/eLife.86439

Schwarzkopf, D. S., Anderson, E. J., de Haas, B., White, S. J., & Rees, G. (2014). Larger Extrastriate Population Receptive Fields in Autism Spectrum Disorders. The Journal of Neuroscience, 34(7), 2713–2724. 10.1523/JNEUROSCI.4416-13.2014

Senden, M., Reithler, J., Gijsen, S., & Goebel, R. (2014). Evaluating Population Receptive Field Estimation Frameworks in Terms of Robustness and Reproducibility. PLOS ONE, S(12), e114054. 10.1371/journal.pone.0114054

Sereno, M. I., Dale, A. M., Reppas, J. B., Kwong, K. K., Belliveau, J. W., Brady, T. J., Rosen, B. R., & Tootell, R. B. (1995). Borders of multiple visual areas in humans revealed by functional magnetic resonance imaging. Science (New York, N.Y.), 268(5212), 889–893.

Sereno, M. I., Sood, M. R., & Huang, R.-S. (2022). Topological Maps and Brain Computations From Low to High. Frontiers in Systems Neuroscience, 16. https://www.frontiersin.org/articles/10.3389/fnsys.2022.787737

Silson, E. H., Reynolds, R. C., Kravitz, D. J., & Baker, C. I. (2018). Differential sampling of visual space in ventral and dorsal early visual cortex. Journal of Neuroscience, 2717–17. 10.1523/JNEUROSCI.2717-17.2018

Stephan, K. E., Weiskopf, N., Drysdale, P. M., Robinson, P. A., & Friston, K. J. (2007). Comparing hemodynamic models with DCM. NeuroImage, 38(3), 387–401. 10.1016/j.neuroimage.2007.07.040

Tangtartharakul, G., Morgan, C. A., Rushton, S. K., & Schwarzkopf, D. S. (2023). Retinotopic connectivity maps of human visual cortex with unconstrained eye movements. Human Brain Mapping, 44(16), 5221–5237. 10.1002/hbm.26446

van Dijk, J. A., de Haas, B., Moutsiana, C., & Schwarzkopf, D. S. (2016). Intersession reliability of population receptive field estimates. NeuroImage, 143, 293–303. 10.1016/j.neuroimage.2016.09.013

Van Essen, D. C., Smith, S. M., Barch, D. M., Behrens, T. E. J., Yacoub, E., Ugurbil, K., & WU-Minn HCP Consortium. (2013). The WU-Minn Human Connectome Project: An overview. NeuroImage, 80, 62–79. 10.1016/j.neuroimage.2013.05.041

Wandell, B. A., Dumoulin, S. O., & Brewer, A. A. (2007). Visual field maps in human cortex. Neuron, 56(2), 366–383. 10.1016/j.neuron.2007.10.012

Zeidman, P., Silson, E. H., Schwarzkopf, D. S., Baker, C. I., & Penny, W. (2017). Bayesian population receptive field modelling. NeuroImage. 10.1016/j.neuroimage.2017.09.008

Zuiderbaan, W., Harvey, B. M., & Dumoulin, S. O. (2012). Modeling center-surround configurations in population receptive fields using fMRI. Journal of Vision, 12(3), 10. 10.1167/12.3.10

