## Supplementary Figures for "Hemodynamic modelling improves population receptive field estimates"

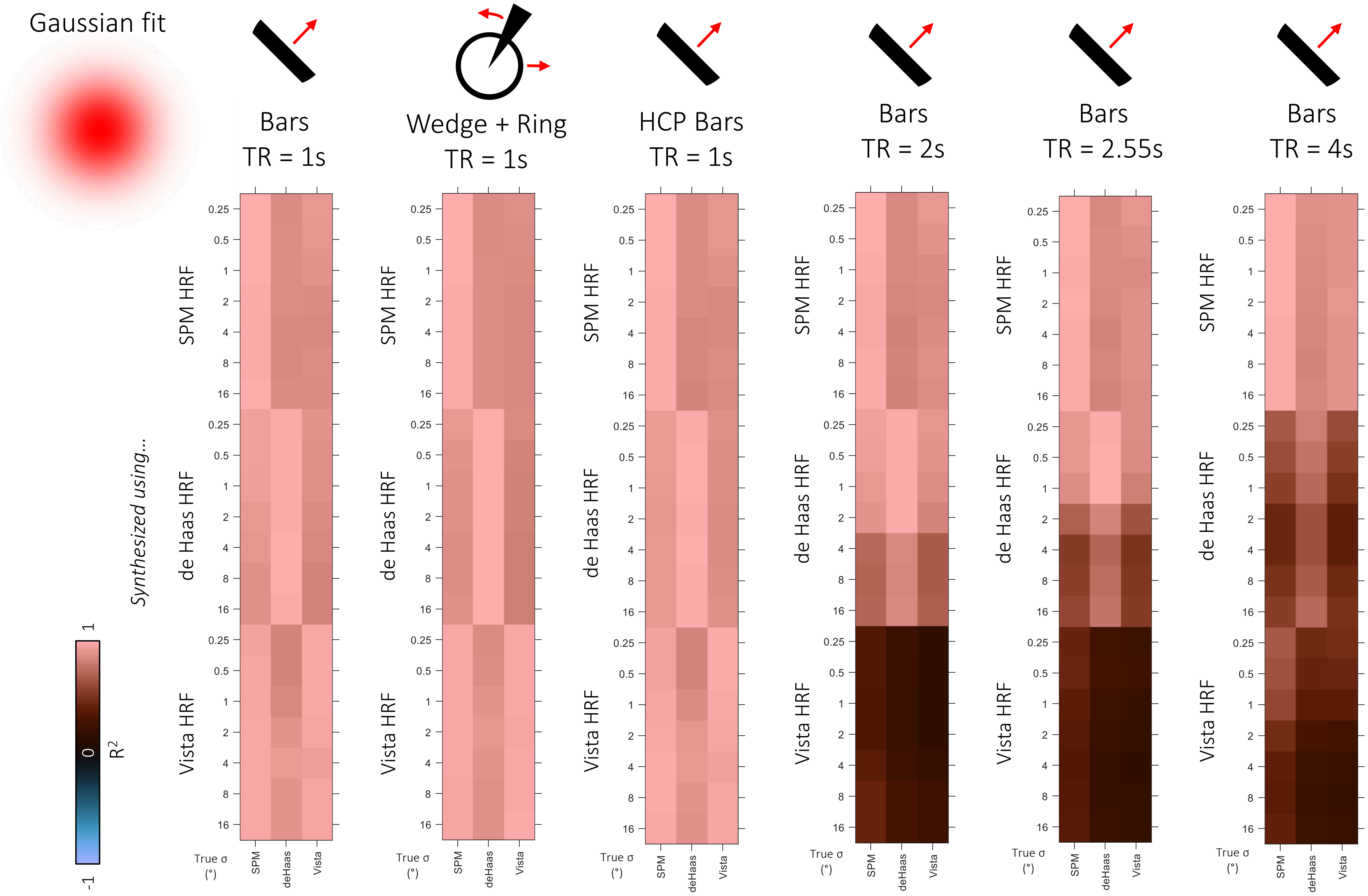


**Figure S1.** Goodness-of-fit ($R^2$) of the average HRF fit for each true pRF size (rows) in each of the three simulated datasets. We fit a standard 2D Gaussian pRF model with concurrent HRF fitting.

The three columns show the HRF fit to each of the three canonical HRFs. Each panel shows a simulation analysis for a different stimulation design (see top row).

Notably, HRF fits are generally good for designs with TR=1 s, but much worse for designs with longer TR, especially for data synthesized with the Vista HRF.

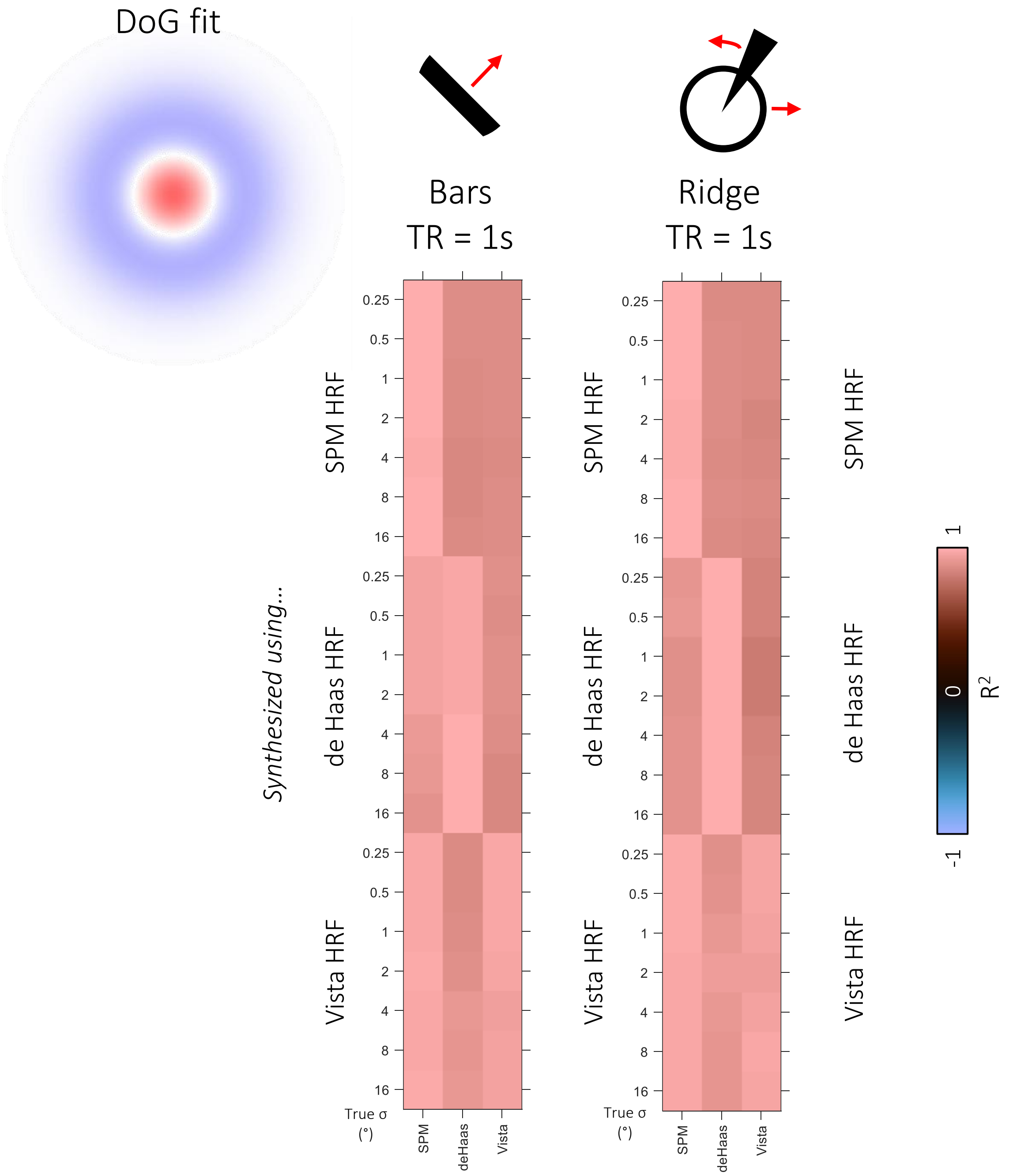


**Figure S2.** Goodness-of-fit ($R^2$) of the average HRF fit for each true pRF size (rows) in each of the three simulated datasets. We fit a center-surround DoG pRF model with concurrent HRF fitting – but note that the true data are based on a 2D Gaussian pRF model.

The three columns show the HRF fit to each of the three canonical HRFs. Each panel shows a simulation analysis for a different stimulation design (see top row).

Notably, HRF fits are generally good for designs with TR=1 s, but much worse for designs with longer TR, especially for data synthesized with the Vista HRF.

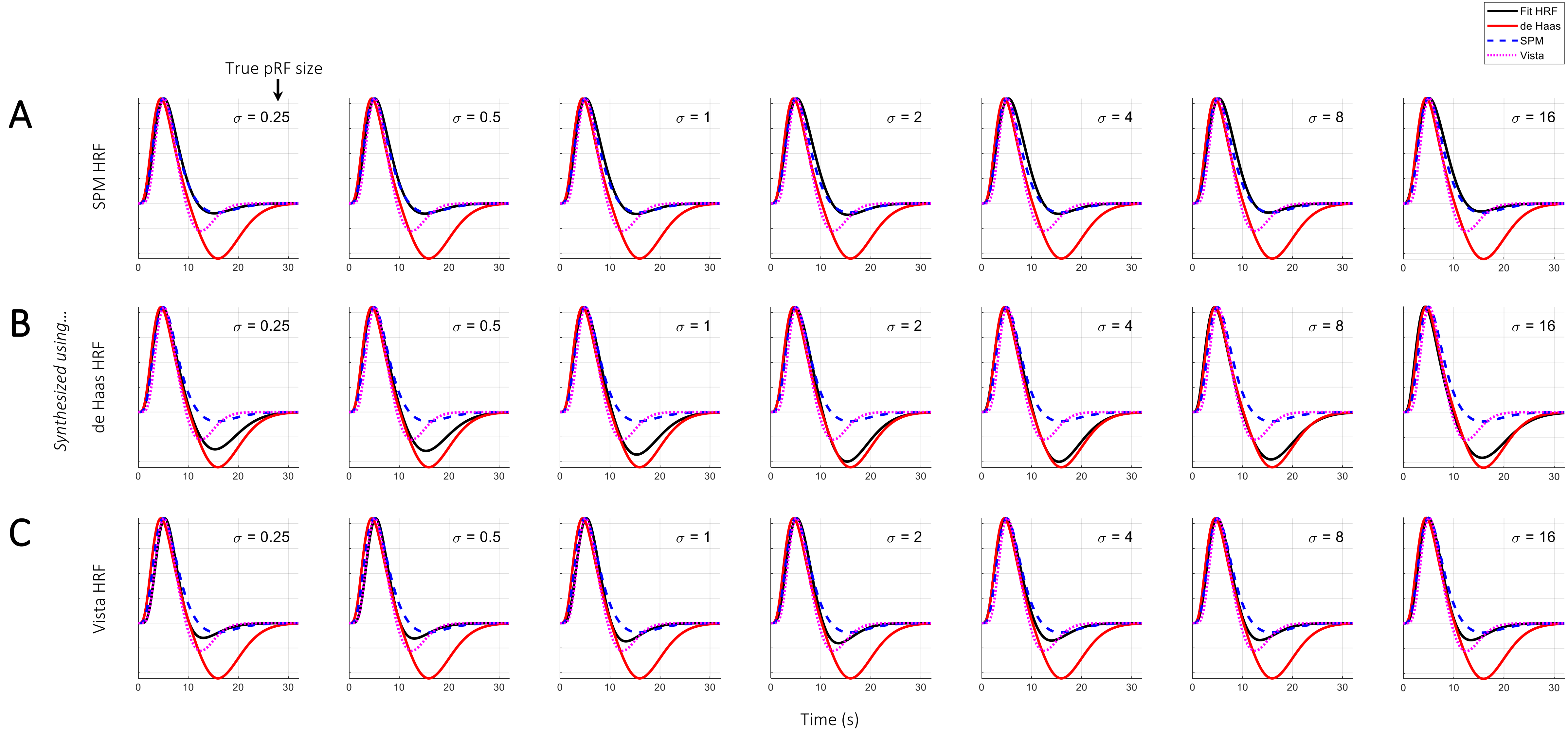


**Figure S3.** Concurrent fit closely estimates the true HRF at TR=1. We synthesized data for a set of biologically plausible Gaussian pRFs but using either the SPM canonical HRF (**A**), our SamSrf de Haas HRF (**B**), or the Vista HRF (**C**). The black line shows the average fit HRF for each dataset from the Concurrent fit a 2D Gaussian pRF model, compared to the three canonical functions (see legend). Columns show the fits for the seven true pRF sizes we simulated.

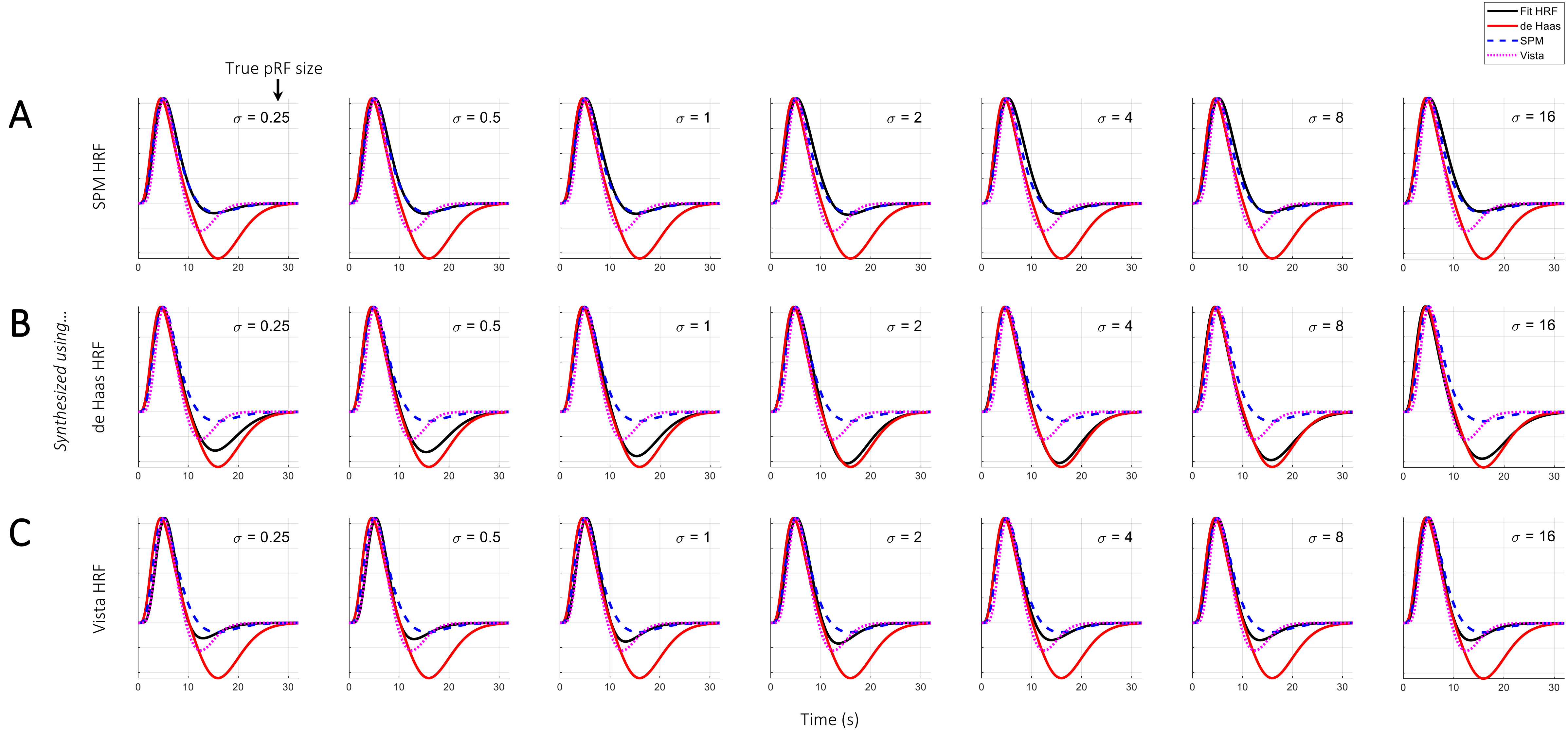


**Figure S4.** Concurrent fit closely estimates the true HRF at TR=1. All conventions as in Figure S3 except that we used a more conservative threshold including only pRF model fits with $R^2>0.2$.

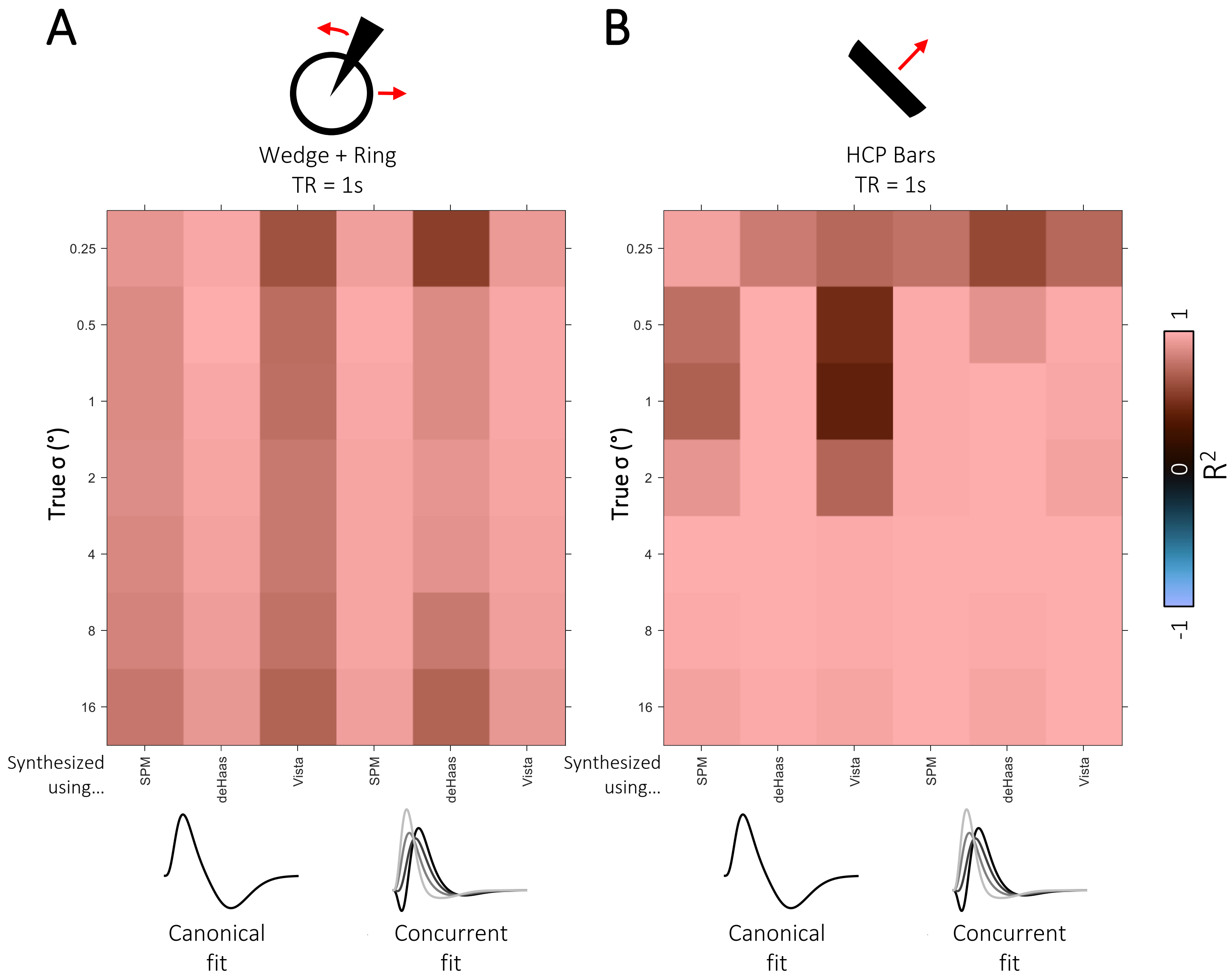


**Figure S5.** Goodness-of-fit between the estimated pRF shape and the true pRF, separately for different true pRF sizes (rows) and the three simulated datasets (columns). The left three columns show Canonical fit results and the right three columns show the Concurrent fit results. Results are shown for data synthesized using a combined wedge-and-ring stimulus (**A**) and a sweeping bar stimulus as used by the HCP (**B**).

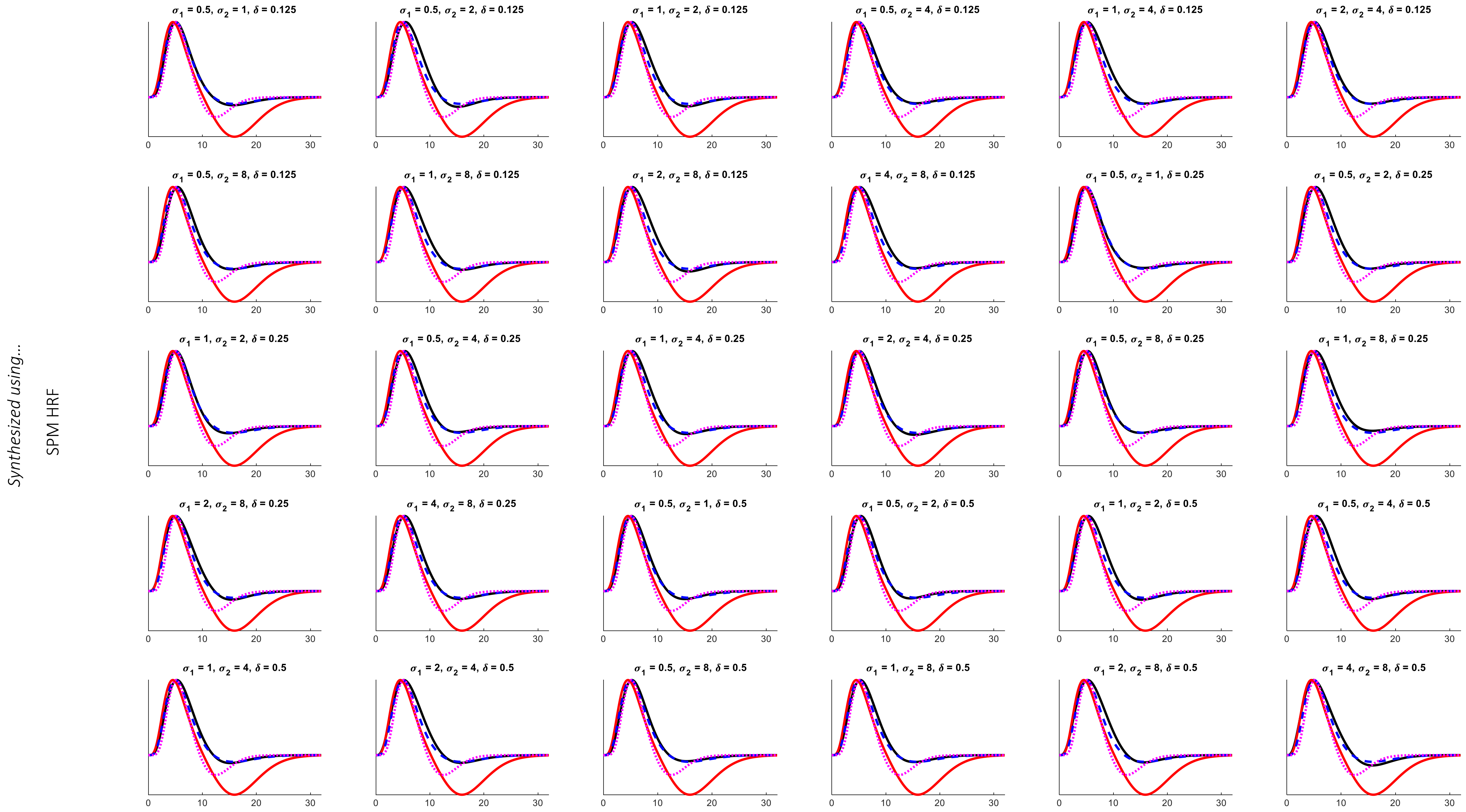


**Figure S6.** Concurrent fit closely estimates the true HRF also for fitting a DoG pRF model. We synthesized data for a set of biologically plausible pRFs based on a DoG center-surround model and the SPM canonical HRF. The black line shows the average fit HRF for each dataset from Concurrent fit of a DoG center-surround model, compared to the three canonical functions (see legend). Each panel shows the fits for a unique combination of true pRF parameters sizes and amplitude ratios we simulated.

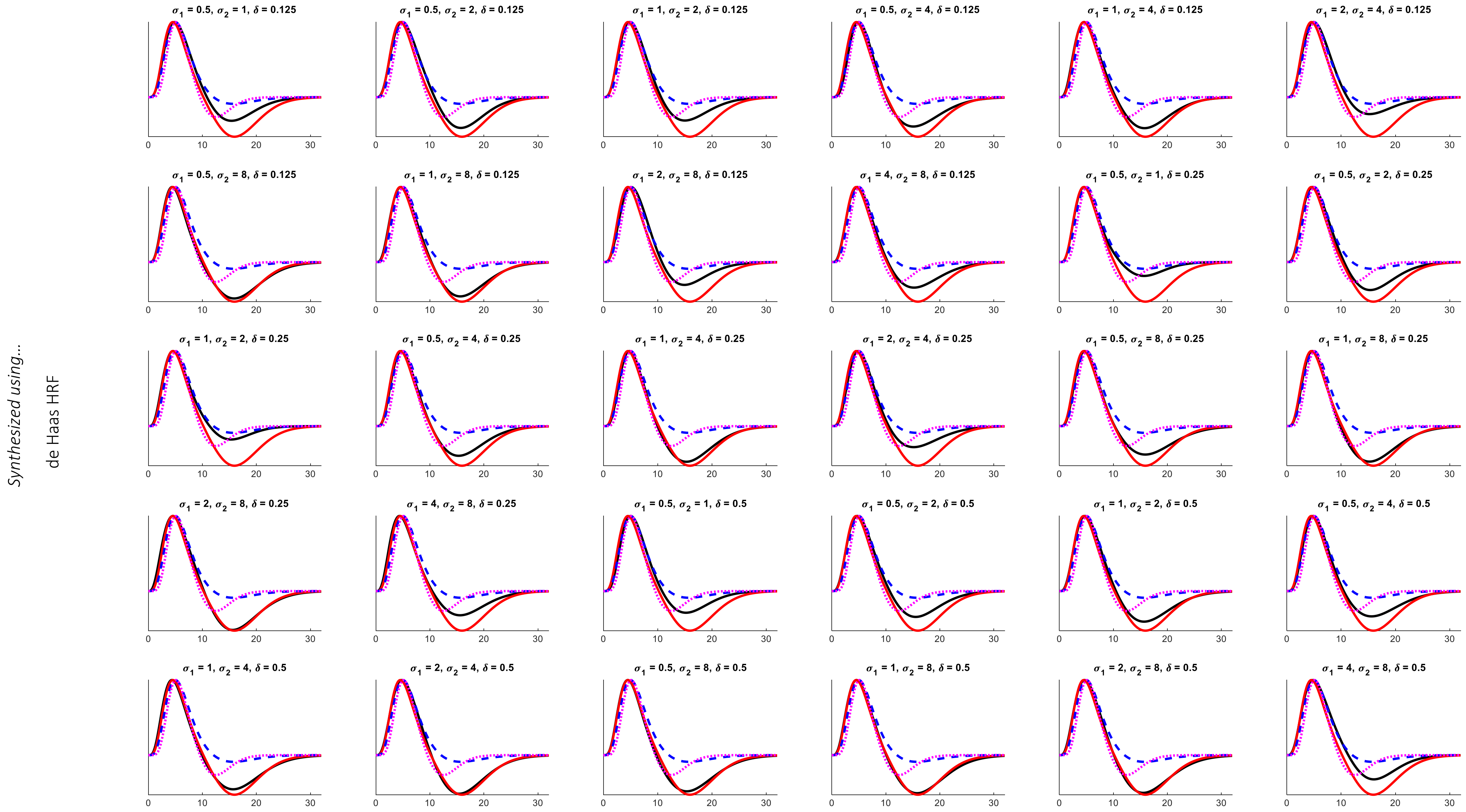


**Figure S7.** Concurrent fit also closely estimates the true HRF also for fitting a DoG pRF model. We synthesized data for a set of biologically plausible pRFs based on a DoG center-surround model and the de Haas canonical HRF. The black line shows the average fit HRF for each dataset from Concurrent fit of a DoG center-surround model, compared to the three canonical functions (see legend). Each panel shows the fits for a unique combination of true pRF parameters sizes and amplitude ratios we simulated.

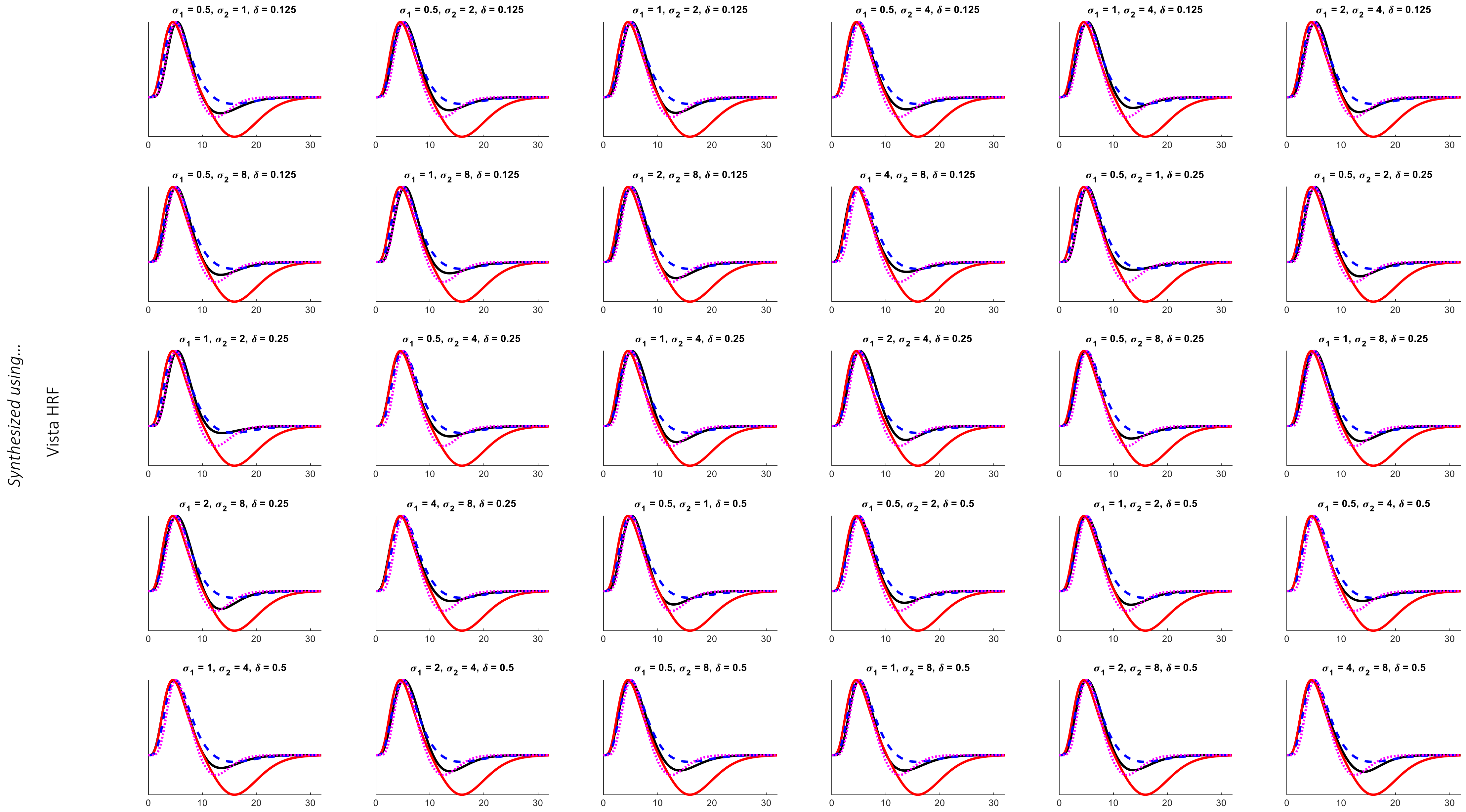


Figure S8. Concurrent fit closely estimates the true HRF also for fitting a DoG pRF model. We synthesized data for a set of biologically plausible pRFs based on a DoG center-surround model and the Vista canonical HRF. The black line shows the average fit HRF for each dataset from Concurrent fit of a DoG center-surround model, compared to the three canonical functions (see legend). Each panel shows the fits for a unique combination of true pRF parameters sizes and amplitude ratios we simulated.

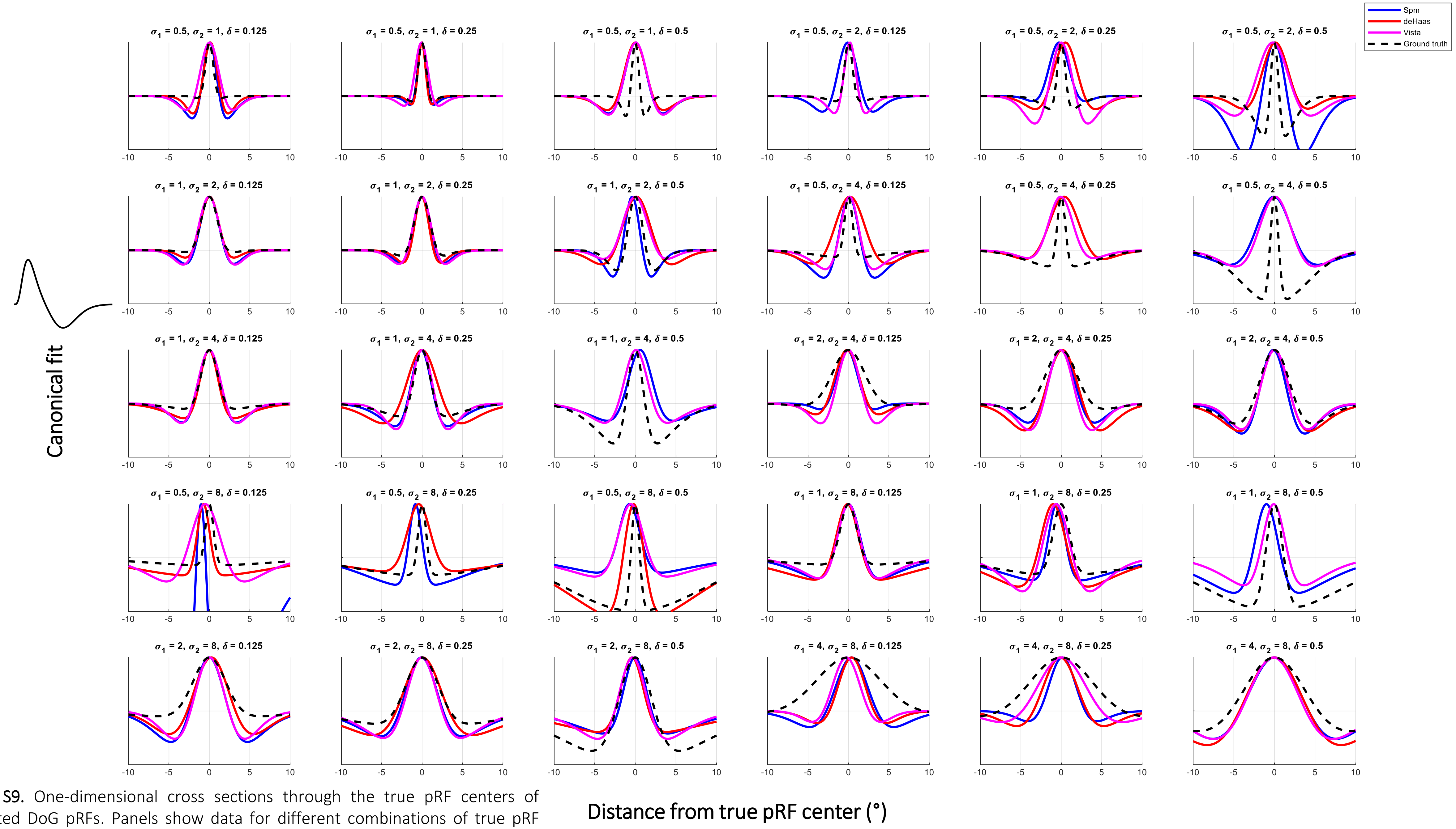


**Figure S9.** One-dimensional cross sections through the true pRF centers of simulated DoG pRFs. Panels show data for different combinations of true pRF parameters. Results from Canonical fit. Other conventions as Figure 3A-D.

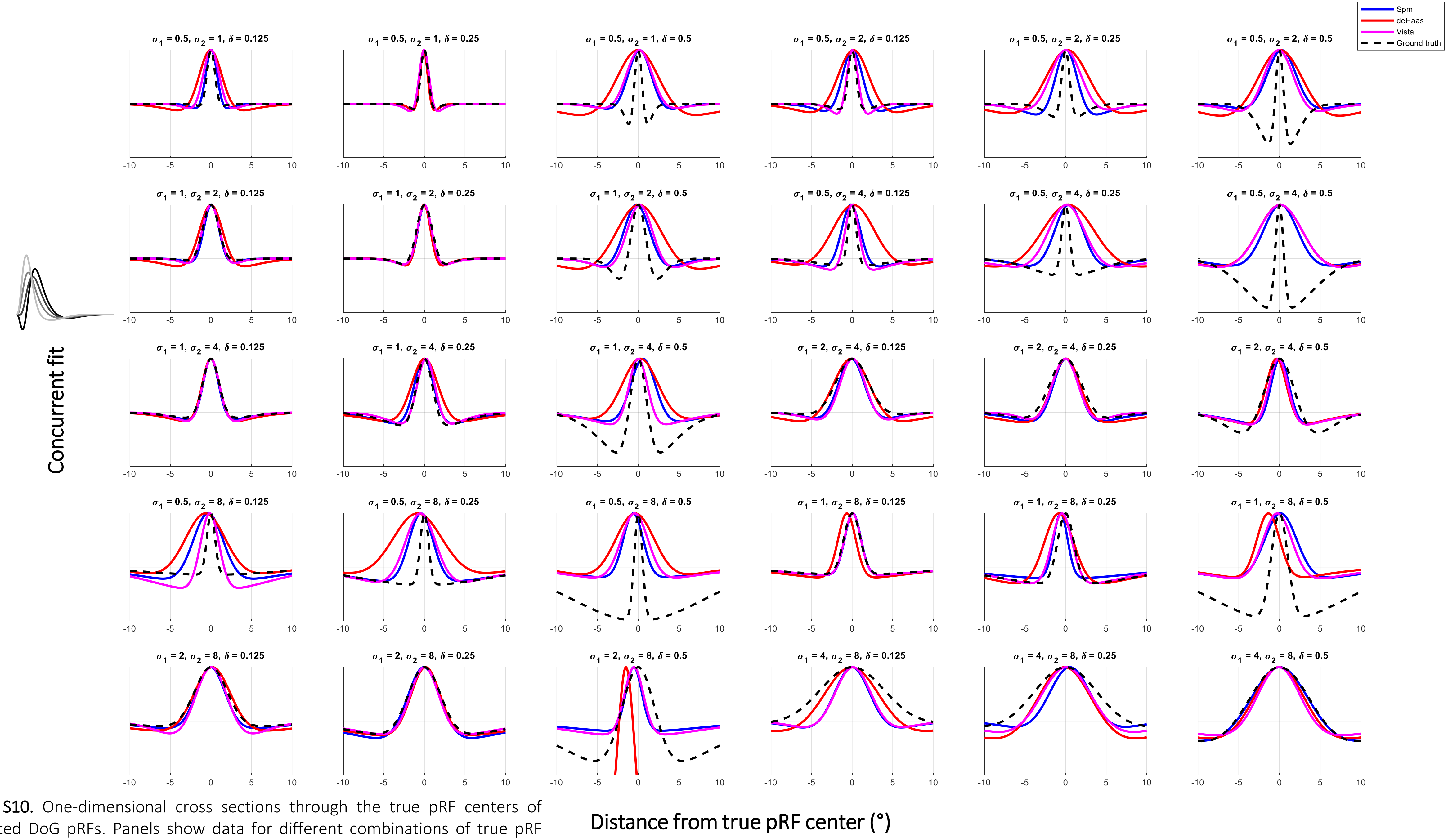


**Figure S10.** One-dimensional cross sections through the true pRF centers of simulated DoG pRFs. Panels show data for different combinations of true pRF parameters. Results from Concurrent fit. Other conventions as Figure 3A-D.

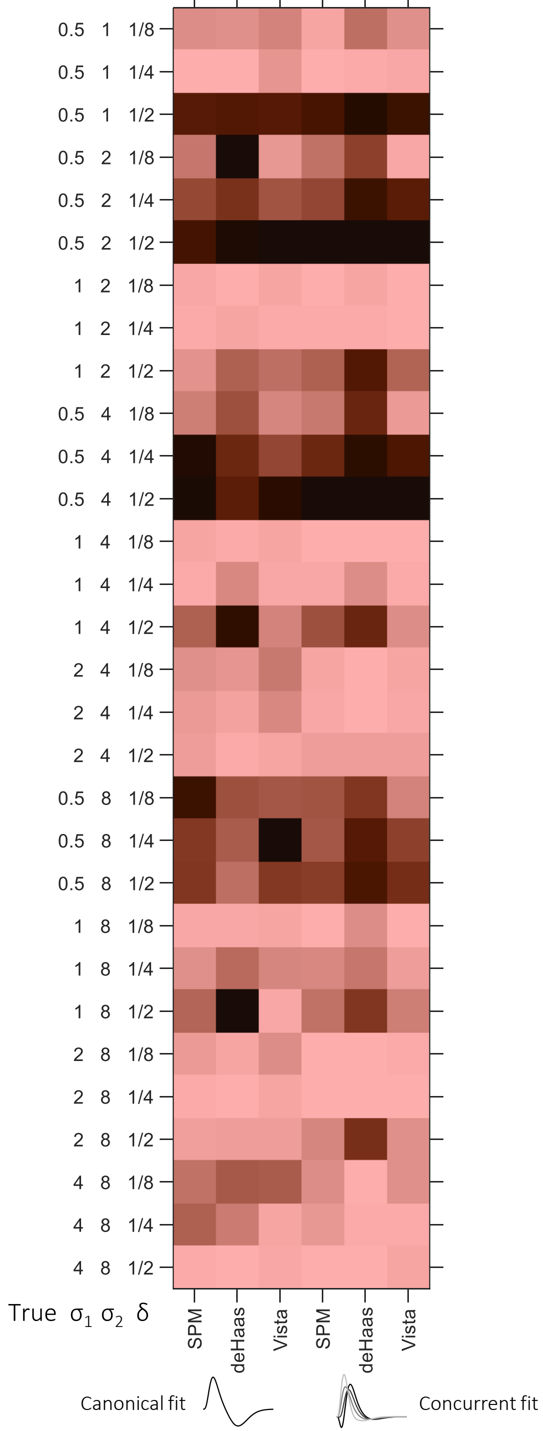


**Figure S11.** Goodness-of-fit between the estimated pRF shape and the true pRF, separately for different combinations of true pRF parameters (rows) and the three simulated datasets (columns). The left three columns show Canonical fit results and the right three columns show the Concurrent fit results. Results are shown for data synthesized using our standard sweeping bar stimulus and a DoG center-surround pRF model.

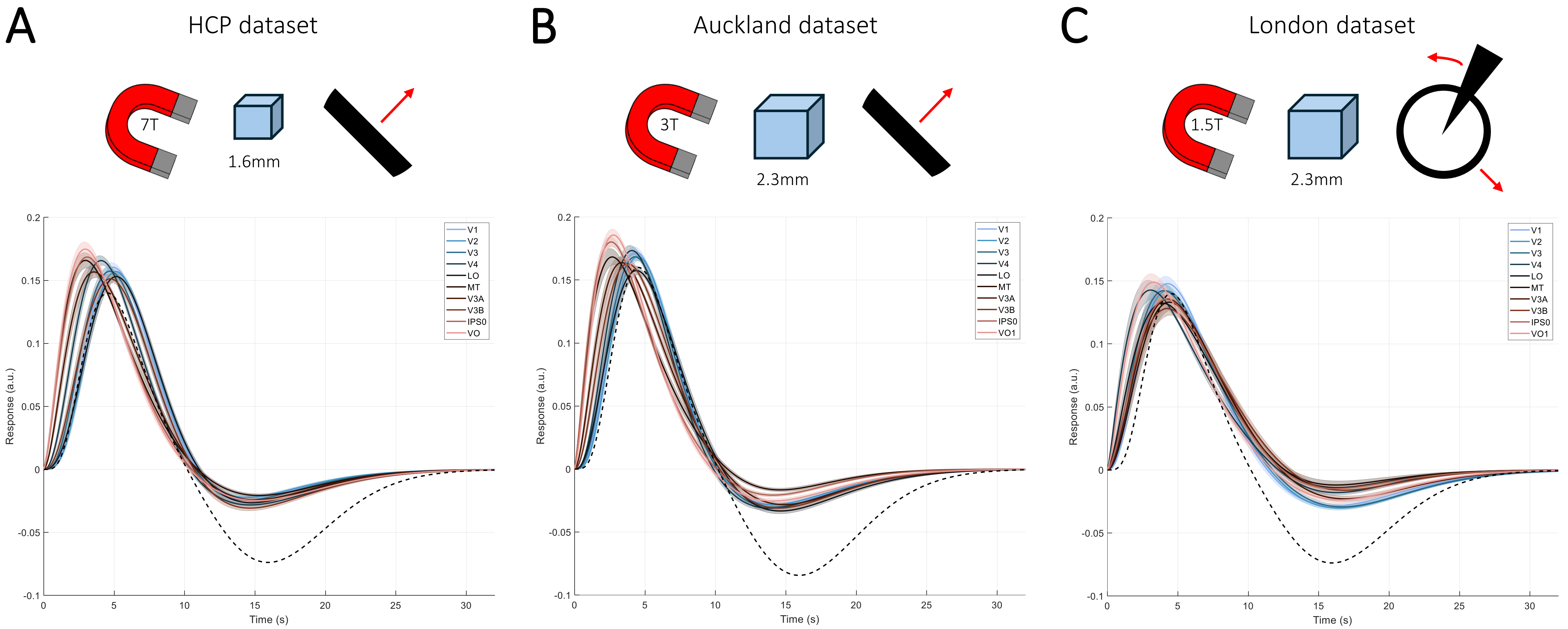


**Figure S12.** Concurrent HRF fitting of empirical pRF data. **A-C**. Average HRFs estimated in Concurrent fit for the HCP (**A**), Auckland (**B**), and London (**C**) datasets. HRFs were averaged separately for several retinotopic areas, and then averaged across participants (shaded regions denote ±1 standard error of the mean across participants). The dashed black line denotes the SamSrf de Haas canonical HRF for comparison.

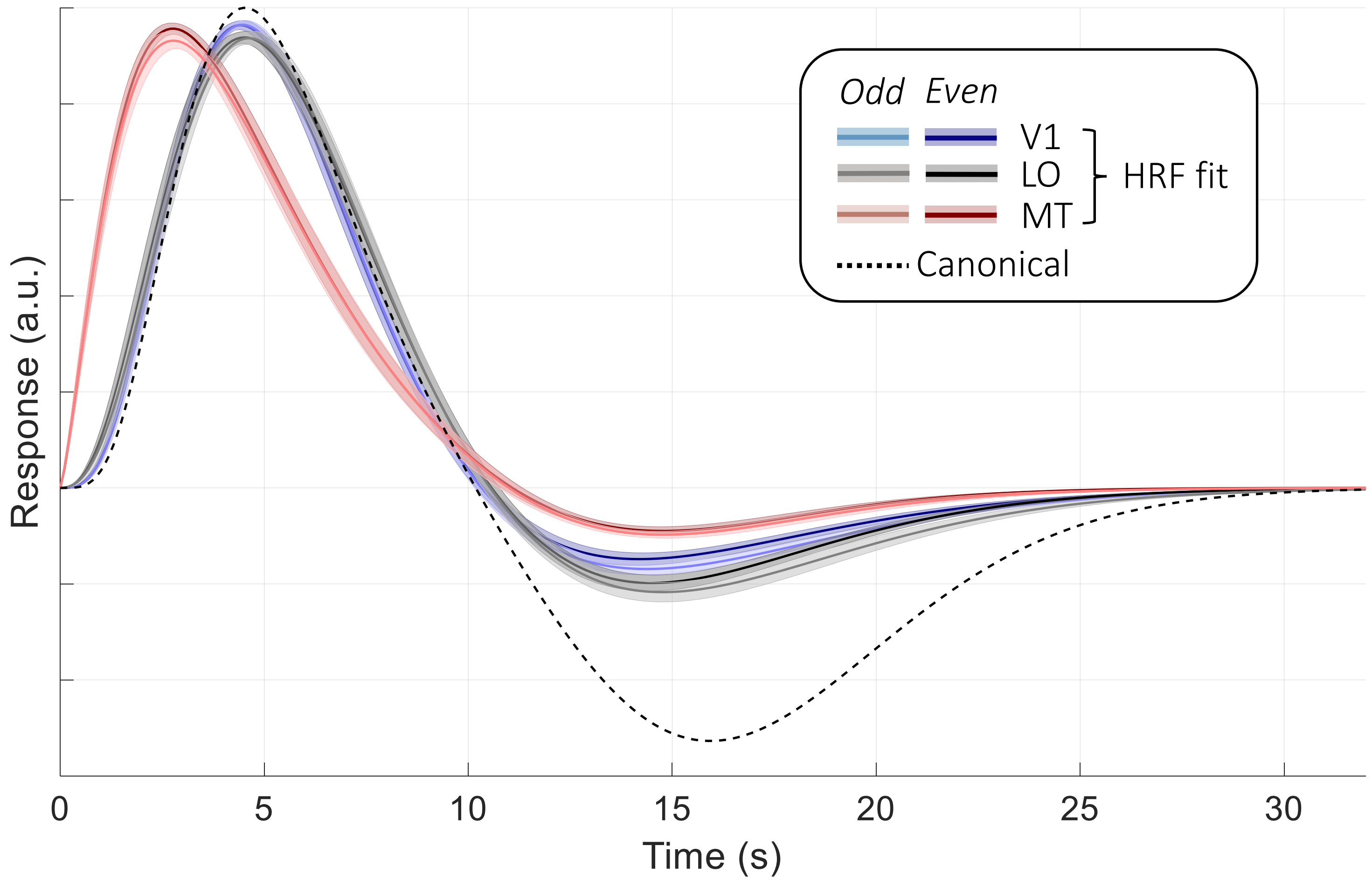


**Figure S13.** Concurrent HRF fitting of cross-validated empirical pRF data from the Auckland dataset. pRF and HRF models were estimated separately for odd- (lighter colors) and even-numbered (darker colors) runs. HRFs were averaged separately for several retinotopic areas, and then averaged across participants (shaded regions denote ±1 standard error of the mean across participants). The dashed black line denotes the SamSrf de Haas canonical HRF for comparison. There is a very close correspondence between the HRFs in a given visual region between cross-validation folds.

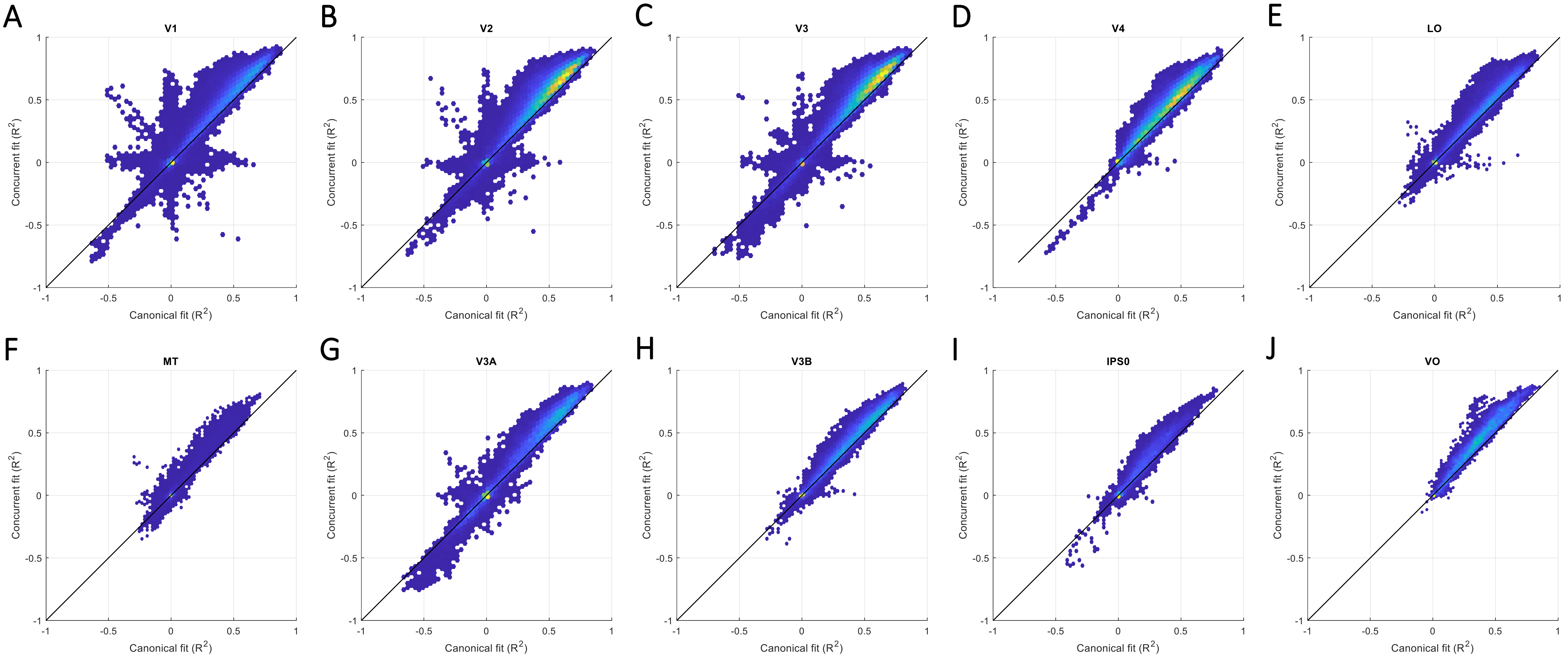


**Figure S14.** Two-dimensional density hex-plots of the cross-validated goodness-of-fit ($R^2$) for the concurrent HRF fit plotted against the corresponding canonical HRF fit. The pRF model was estimated separately for odd- and even-numbered runs. Cross-validated $R^2$ measures how well the predicted pRF (and HRF, in concurrent fit) model fits the observed data from the independent runs. These $R^2$ values were averaged across both cross-validation folds. Cross-validated $R^2$ can be negative for very poor fits. Data shown are pooled across all 23 participants. It is evident that the concurrent fit is generally better than the canonical fit, at least for vertices with generally positive $R^2$. Data are separated by visual region V1 (**A**), V2 (**B**), V3 (**C**), V4 (**D**), LO (**E**), MT (**F**), V3A (**G**), V3B (**H**), IPS0 (**I**), and VO (**J**).
